# Defining Epidermal Stem Cell Fate Infidelity and Immunogenicity in Hidradenitis Suppurativa at the Single-Cell Resolution

**DOI:** 10.1101/2020.04.21.053611

**Authors:** Meaghan Marohn, Meng-ju Lin, Wei-wen Yu, Ciara Mae Mendoza, Juliana Remark, Alireza Khodadadi-Jamayran, Ernest S. Chiu, Catherine Pei-ju Lu

## Abstract

Hidradenitis suppurativa (HS) is a severe chronic inflammatory skin disease affecting human apocrine sweat gland-bearing skin regions. One unique feature of HS is the development of keratinized sinus tracts that grow extensively deep in the dermis and are highly immunogenic. Here, we demonstrated that the stem cell fate infidelity exists in the HS sinus tracts, which exhibit features of both surface epidermis and appendages. Using single cell transcriptome analyses, we finely dissected different compartments of the HS epithelium and identified their respective changes in cytokine expression during disease progression and the critical interactions with the immune cells. Together, our work provides advanced understanding of the pathological epidermal remodeling and important implications for HS therapeutics.

## Introduction

Hidradenitis suppurativa (HS) is a chronic inflammatory skin disease with acutely painful flares primarily affecting human apocrine sweat gland-bearing skin regions, including axillae, perineum, groin, and inframammary regions^1,2^. The psychosocial burden of the disease is particularly destructive because of extreme pain, malodorous discharge, and intensive scarring in these skin areas^3^. The overall prevalence of HS ranges from 0.05-4.1% with higher occurrence among females and African Americans, and strong associations with smoking and obesity^4^. Due to incomplete understanding of its pathogenesis, the treatment and management of HS in patients has been extremely difficult. Combined medical and surgical treatments are recommended according to disease severity; however, wide excisional surgical procedures are often required for advanced or recurrent lesions^5^. The disease has a bimodal distribution of onset, once during puberty when the skin appendages (hair follicles and sweat glands) in these areas become mature and functional and again in women around 40 years likely due to hormonal changes perimenopausally.

One unique feature that distinguishes HS from other inflammatory skin diseases is the extensive formation of sinus tracts (also known as fistulae or tunnels^6^) which develop deep into the lesional dermis as disease progresses. The distinct inflammatory components found surrounding HS sinus tracts recently suggest that these structures may be potential and active drivers of the chronic inflammation^7,8^. Additionally, the disappearance of skin appendages (pilosebaceous-apocrine units) in the HS lesions has been frequently observed ^9^, which prompts speculations on the cells of origin for HS sinus tracts and the pathological mechanisms that trigger their formation.

Previous works using mouse models have demonstrated that adult stem cells and progenitor cells exist in different part of the skin epidermis and appendages^10–13^ and that important transcriptional regulations define their cell fates^14^. During normal development and homeostasis, key transcription factors are expressed in different skin stem cells and establish specific gene expression patterns to maintain their lineage cell fate identity. Loss of these master transcription factors often leads to transformation to a different cell fate. For example, loss of Lhx2 in hair follicle (HF) stem cells causes a switch from HF fate to sebaceous gland fate ^15^; loss of Sox9 in the HF causes a switch from HF fate to interfollicular epidermal cell fate upon activation^16^. Further, it was demonstrated that stem cells exhibit infidelity in wound repair and cancer progression with a feature of co-expression of Sox9 (appendage stem cell fate) and Klf5 (epidermal stem cell fate), and that this cell fate infidelity/duality has functional implications for tumor maintenance^17^. Hidradenitis suppurativa, as a severe chronic inflammation with extensive tissue remodeling and expansion, has been hypothesized to resemble wound and tumor conditions, and yet exhibits its unique feature of developing extensive sinus tracts.

To understand the roles of sinus tracts and skin appendages in HS pathogenesis, we investigated into the stem cell fate commitment and employed single cell RNA-sequencing technology to define transcriptional changes in different lineages of the skin epidermis of HS perilesional and lesional skin. We demonstrated that cells in HS sinus tracts, although similar to interfollicular (surface) epidermis in morphology, express genes specific to sweat and sebaceous glands, and exhibit a significant level of stem cell fate duality. Our computational analyses for scRNAseq data provided insights into the lineage relationships at the transcriptome level and showed that the cells in HS sinus tracts are closest to those in infundibulum of pilosebaceous-apocrine unit. Further, we analyzed the critical cytokines and receptors expressed in the HS samples, and established the immune-epithelium interaction network that contribute to different stages in HS pathogenesis. Our work provides novel insights into epidermal stem cell identity in the context of a human skin disorder with severe chronic inflammation, as well as advances in understanding on how perturbed epidermal cells trigger and interact with the immune system, directly impacting on HS management and therapeutics.

## Result

### Epidermal abnormality occurs prior to immune cell recruitment

The specific skin regions where HS lesions develop (mostly apocrine sweat gland-bearing regions) argue against a general immune dysregulation in the skin. We sought to understand the unique role of the pilosebaceous-apocrine unit in the HS pathogenesis, specifically the epidermal abnormality and immunogenicity. We collected lesional and perilesional skin samples from HS patients^18^ and performed multiplexed immunofluorescent imaging for various immune cell markers. We showed that albeit the normal appearance on the skin surface, perilesional epidermis exhibits significant psoriasiform hyperplasia, while very little immune cell infiltrates into this area (sup figure 1 A-D). Quantification of numbers of immune cells at representative HS perilesional and lesional skin were performed, shown in sup figure 1E. We detected an increased number of macrophages in the lesional perifollicular areas, an increased number of all T cell, B cell and macrophage in lesional epidermis, and the highest level of immune cell infiltration in HS sinus tracts. These results indicate that epidermal abnormality occurs before immune cell recruitment in HS and that aberrant epidermal cells in HS sinus tracts are highly immunogenic, which presents a unique feature of HS pathology amongst all inflammatory skin disorders.

### Keratinocytes lining HS sinus tracts express high levels of sebaceous and sweat gland-specific marker genes

It has been previously documented that infundibular HF occlusion and loss of appendages are frequently observed in HS lesions ^19–21^. We posited that HS lesional sinus tracts may be derived from skin appendages and that cells lining the sinus tracts may still express reminiscent signature genes of the pilosebaceous-apocrine units. While we did not detect the expression of LHX2, an HF specific marker, in HS sinus tracts (Sup Figure 2), we showed significant expression of SCD1 (specific for normal sebaceous gland and apocrine sweat gland) (Figure 1A) and CEA (specific for normal eccrine sweat gland) (Figure 1B) in distinct layers of stratified keratinocytes lining the HS lesional sinus tracts. We examined samples from various HS patients and aimed to determine whether expressions of these markers is homogeneous. Shown in Figure 1C, the keratinocytes in HS sinus tracts exhibit a striking heterogeneity with regards to SCD1 and CEA expression, suggesting that the cells of origin in HS sinus tracts could be epidermal stem cells that possess multipotency to give rise to cells with various glandular features. In addition, we showed that the expression of SOX10, a marker for sweat gland tumors^22^, is upregulated in HS lesional sweat glands and the outermost suprabasal layer of sinus tracts (Figure 1D). Most interestingly, nuclear localization of androgen receptor (AR), exclusively detected in sebaceous and apocrine sweat glands in normal skin, is now detected at a high level in HS sinus tracts (Figure 1E), which also highlight the potential involvement of hormones in the HS onset. Altogether, these results support the notion that cells lining sinus tracts are derived from progenitors that have the potential to give rise to sweat and sebaceous glands.

**Figure 1.**
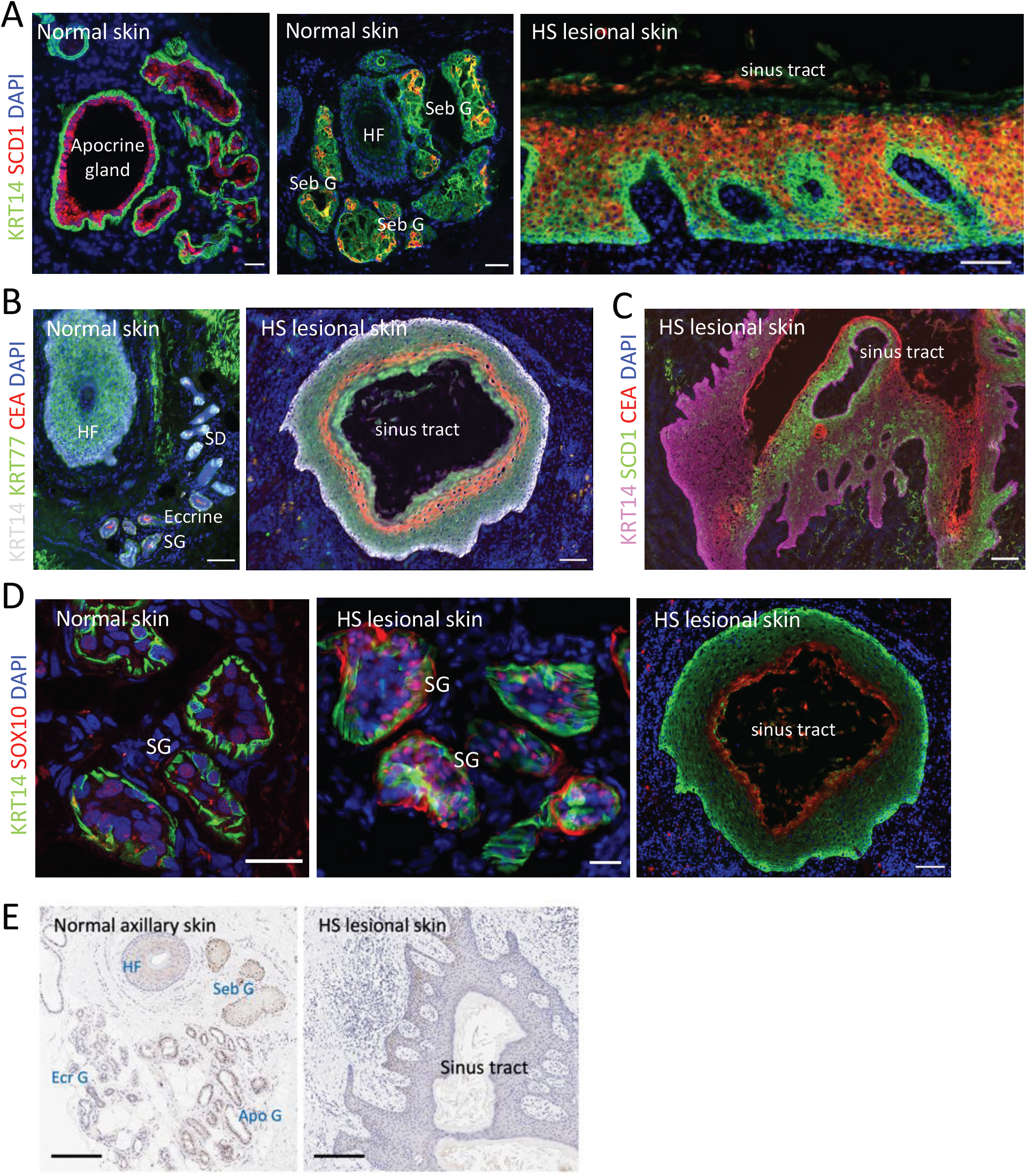
Keratinocytes lining HS sinus tracts express high levels of sebaceous and sweat gland-specific marker genes. (A) Immunofluorescent images of normal axillary skin and HS lesional skin, showing expression of SCD1 in red. Note that SCD1 is highly expressed in the luminal cells of the apocrine gland, sebaceous gland (Seb G) (not in hair follicle HF), as well as HS sinus tract. Scale bar (left to right), 25 μm, 50 μm, 100 μm, respectively. (B) Immunofluorescent images of normal and HS lesional skin, showing staining of KRT77 in green and CEA in red. Note that KRT77 is expressed in normal HF and sweat duct (SD). CEA is expressed in normal eccrine sweat gland (SG). Both of them are expressed in the HS sinus tract. Scale bar, both 100 μm. (C) Immunofluorescent images of HS lesional skin, showing expression of SCD1 and CEA in HS sinus tract. Note that these two markers are expressed in distinct layers within the sinus tract epithelium. Scale bar, 200 μm. (D) Immunofluorescent images of normal skin and HS lesional skin, showing staining of SOX10 in red. Note that SOX10 is expressed at a low level in normal sweat gland and high level in HS lesional sweat gland. It is also present in outer most layer of HS sinus tract. Scale bar (left to right), 25 μm, 25 μm, 100 μm, respectively. (E) Immunohistochemical staining of androgen receptor (AR) in normal axillary skin (left) and HS lesional skin (right). Scale bars, 200 μm. Note that AR is expressed in normal sebaceous gland and apocrine gland (not in HF, nor eccrine SG), and also extensively in the HS sinus tract.

### Epidermal Stem cell fate duality occurs in HS sinus tracts

One possibility to explain the formation of sinus tracts and loss of appendages in the HS lesions is that the cells in the appendages lost the key transcription factors governing appendage fate and turned on the interfollicular epidermal differentiation programs as exhibited in the HS sinus tracts, such as KRT10 expression shown in sup Figure 3. We sought to investigate whether stem cell fate infidelity (or duality, exhibiting both interfollicular epidermal fate and appendage fate) exists in sinus tract formation in humans and contributes to HS pathogenesis. Different from mouse skin appendages that are strictly Sox9+ (very little Klf5 or Krt10 expression), the upper part of human appendages (hair follicle infundibulum and sweat ducts) has extensive co-expression of SOX9 and KLF5 (80%), as well as high level of KRT10 during normal conditions (Figure 2A-D and sup Figure 3), suggesting that cells in this area already exhibit significant stem cell fate duality during normal homeostasis. Further, we found that while mouse interfollicular epidermis exclusively expresses Klf5 ^17^, the majority of the human epidermal keratinocytes expresses KLF5 with few cells (4%) expressing SOX9 in the basal layer (Figure 2A). Interestingly, in HS lesional skin, more cells in surface interfollicular epidermis (8%) express SOX9 and frequently at the bottom protrusion of the hyperplastic ridges (Figure 2E). Nevertheless, cells in the interfollicular epidermis from both normal and HS skin exhibit distinct fates, with little co-expression of KLF5 and SOX9. Importantly, in HS sinus tracts, we detected a significant percentage of SOX9+ cells, as well as SOX9+KLF5+ cells (Figure F-H). Quantification in Figure 2I shows that 70% of the basal cells are SOX9+, amongst which 40% are also KLF5+; at the protrusion front of the sinus tracts, about 23.5% of the cells are SOX9+ and amongst which 55% are also KLF5+. Here, we demonstrated that stem cell fate infidelity/duality occurs at a high frequency in the cells of HS sinus tract cells, at a level between HF infundibulum/SD area and interfollicular epidermis. The distribution of epidermal cells exhibiting distinct or dual cell fate in normal and HS skin are illustrated in Figure 2J.

**Figure 2.**
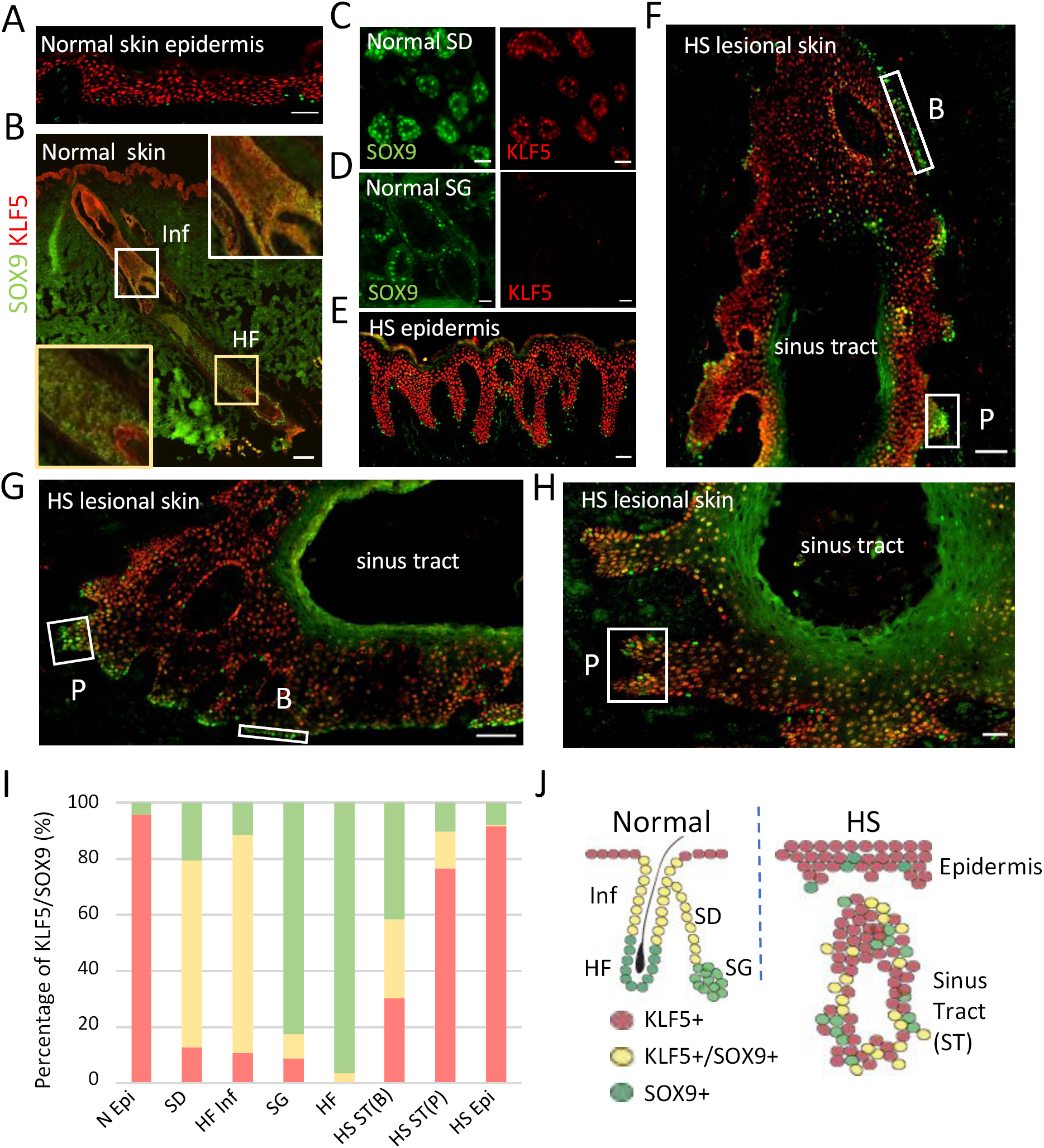
Epidermal stem cell fate duality occurs in the HS sinus tracts. (A-F) Immunofluorescent images of normal skin (A-D) and HS lesional skin (E-H), showing expression of KLF5 (interfollicular epidermal fate) in red and SOX9 (appendage fate) in green. Note that KLF5 is expressed dominantly in epidermis of normal and HS skin and SOX9 restricted to lower part of HF and SG, while both are expressed in upper part of HF (Infundibulum, Inf) and sweat duct (SD) and significant portion in the HS sinus tract. Scale bar, (A) 50 μm, (B) 200 μm, (C-D) 25 μm, (E) 50 μm, (F-G) 100 μm and (H) 50 μm. Boxes in F-H, showing representative areas for sinus tract basal layer (B) and protrusions (P). (I) Quantification of cell numbers in different skin regions, showing percentage of KLF5+(red), KLF5+/SOX9+ (yellow), and SOX9+ (green) cells. n (from left to right) = 5, 12, 10, 23, 10, 11, 11 and 8. (J) Diagram illustrating the distribution of cells in normal and HS skin with surface epidermal fate (KLF5+), appendage fate (SOX9+), and dual fate (KLF5+/SOX9+).

### Single cell RNAseq reveals changes in the epidermis and skin appendages in HS pathogenesis

To understand the cells of origin that give rise to HS sinus tracts, we employed single cell RNA sequencing to acquire data on a total of 5,848 keratinocytes from normal, HS perilesional and lesional skin of axillary and groin regions. As shown in the UMAP plot in Figure 3A, the keratinocytes were clustered into 17 groups using the Seurat v3.0 workflow^23,24^. A heatmap of the top genes in each cluster as well as a Dot Plot of known signature genes for different cell types in the skin epidermal lineages were generated to determine cell identity (Figure 3B and sup Figure 4). We were able to identify distinct clusters for different types of appendages, hair follicle, sebaceous glands, sweat gland myoepithelial cells, eccrine luminal cells, apocrine luminal cells. Within each of these, cells from different patients (both normal and HS individuals) were clustered into one population. Interestingly, interfollicular epidermis from normal individuals were clustered into one, but that from different patients were separated into different clusters. We performed analyses to look for top genes in these epidermal clusters from different patients and found significant heterogeneity amongst different patients; for example, CCL20 expression is high in patient Ax1, lower in patient Gr1, and absent in others (sup Figure 5).

**Figure 3.**
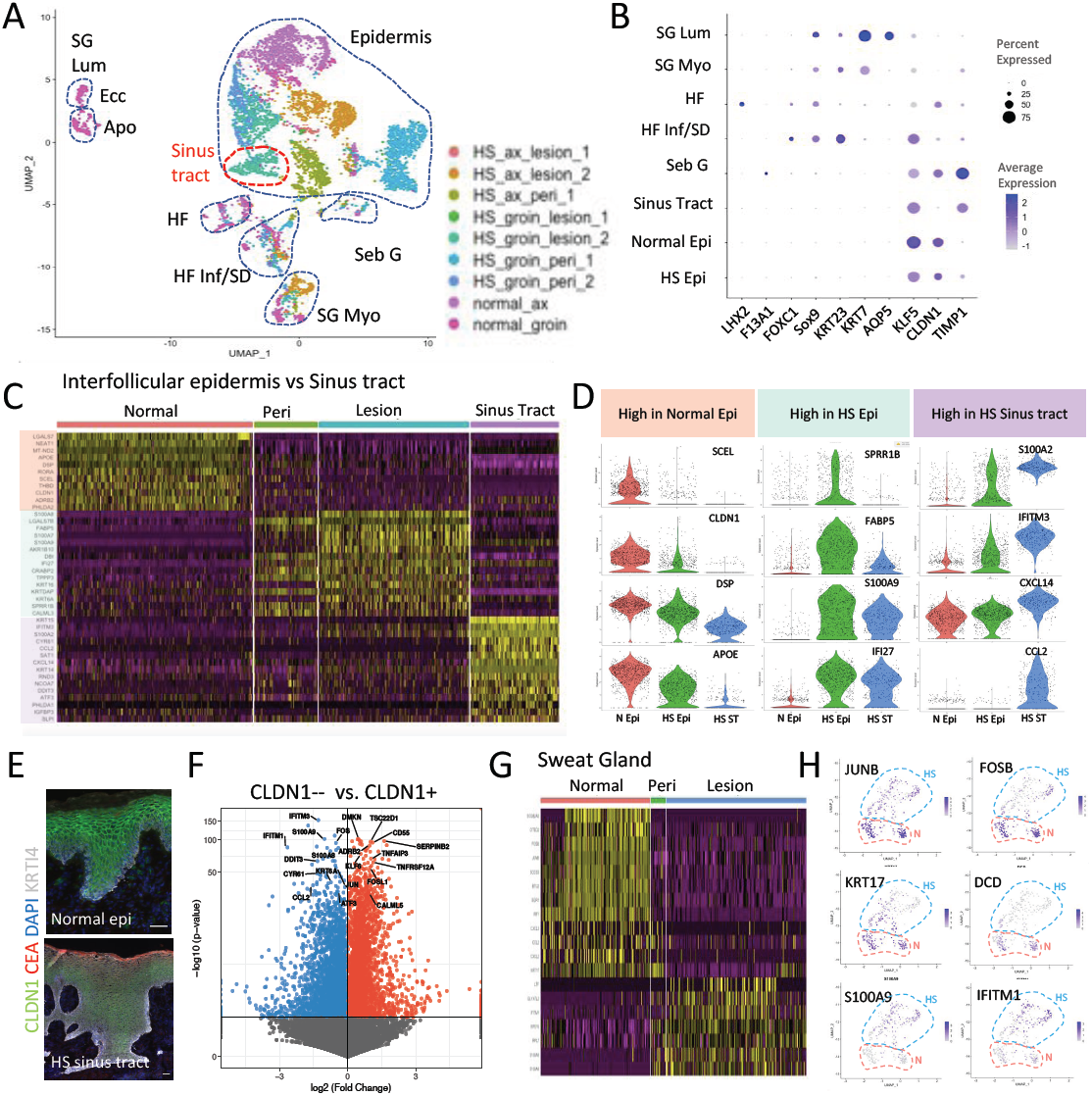
Single cell RNAseq reveals changes in the epidermis and skin appendages in HS pathogenesis. (A) UMAP, unsupervised clustering of keratinocytes from nine different skin samples. Dashed lines circle the clusters that share the same cell identity. (B) Dot plot, showing the percent of expression and average expression level of key genes in each cluster. (C) Heatmap, showing top 10 genes from normal epidermis, HS lesional epidermis, and HS sinus tract. Note that majority of the top genes in the lesional epidermis is also expressed in the perilesional epidermis of the same patient. (D) Violin plot, showing selective genes that are expressed at a high level in normal epidermis, HS epidermis and HS sinus tract (ST). (E) Immunofluorescent images of normal and HS epidermis, showing CLDN1 expression in green and CEA in red. Note that CLDN1 expression is high in normal epidermis, but variable in the HS sinus tract: (left) representative region with high CLDN1 and no CEA expression; (right) representative region with downregulated CLDN1 expression and significant CEA expression. Scale bar, all 100 μm. (F) Volcano plot, showing genes that are differentially expressed in CLDN1+ versus CLDN1-epidermal cells in HS skin. (G) Heatmap, showing top genes expressed in normal v.s. HS lesional sweat gland myoepithelial cells. (H) Feature plot, showing changes in the expression of JUNB, FOSB, KRT17, DCD, S100A9, IFITM1 in normal (red dash-lined) v.s. HS lesional (blue dash-lined) sweat gland cells.

Next, we analyzed the top genes from interfollicular epidermis (of normal, perilesional and lesional), in comparison with HS sinus tracts, and found that many of the most striking changes in lesional epidermal tissue had already preluded in perilesional skin (where little immune cells are recruited) (Figure 3C and D), including genes involved in stress response (S100A7, S100A8, S100A9, SPRR1B, KRTDAP), interferon response (IFI27), retinoic acid regulation (LGALS7B and CRABP2) and lipid metabolism (DBI and FABP5). Secondly, we identified genes that are differentially expressed in HS sinus tracts, such as upregulation of CCL2 and CXCL14 (important for recruitment of macrophages and dendritic cells), CYR61, IGFBP3 and RND3 (TGFB response), ATF3 and DDIT3 (stress response), and IFITM3 (interferon response). We also detected a downregulation of CLDN1 in sinus tract in scRNAseq data and performed an immunofluorescent staining for CLDN1. As shown in Figure 3E, CLDN1 is highly expressed in normal epidermis, while in HS sinus tract, we detected variable levels of CLDN1 expression. Intriguingly, in the region where CEA is expressed, CLDN1 expression is significantly downregulated (also in Sup Figure 6). We performed differential expression analyses for CLDN1+ v.s. CLDN1-cells in the HS epidermis and found that CCL2, S100A8 and S100A9, as well as other stress response genes are upregulated when CLDN1 is not present (Figure 3F). Together with decreased expression of SCEL and DSP, these results emphasizes the importance of barrier function in the HS pathogenesis.

Further, we investigated into the changes in gene expression within the sweat gland myoepithelial cells in normal, perilesional and lesional skin, and detected downregulation of AP1 family members (JUNB and FOSB), important stem cell marker KRT17 and antimicrobial peptide DCD, as well as upregulation of S100A8, S100A9 (stress response) and IFITM1 (interferon response) (Figure 3G-H). Changes in HF during disease progression include increased expression of S100A9, SOX4, CXCL14, KRT6B, and KRT16, and decreased expression of JUNB and FOSB (sup Figure 7). Note that downregulation of AP-1 transcription factor family members (JUNB, FOSB) are detected in both hair follicles and sweat glands in HS skin, suggesting that AP-1 members may play a role in skin appendages during homeostasis, and that loss of AP-1 activity may lead to production of cytokines to recruit neutrophils and macrophages to the epidermis, consistent with works previously demonstrated in psoriasis model^25^.

### HS sinus tracts share the closest lineage relationship with infundibulum of pilosebaceous-apocrine unit

To understand the lineage relationships between HS lesional epidermis, sinus tracts, and appendage clusters, we first performed correlation matrix analysis. As shown in Figure 4A, we found that the sinus tract cluster is grouped with all the clusters from skin appendages and away from interfollicular epidermal clusters. Further, using pseudotime analysis by Monocle v3.0^26^, we showed in Figure 4B that normal epidermal cells are distributed at one middle stem, and branch into lesional epidermis on the right and appendage clusters on the left, while cells from HS sinus tracts split into two cell states aligned with the branches of either epidermis or appendages. Using iCellR interactive tool^27^, we investigated the cell composition of different cell states (I and II) consisting of HS sinus tract cells. As shown in Figure 4B, state I contains predominantly 85% of total sinus tract cells, which share the same state with cells of infundibulum/sweat duct and hair follicles; while state II contains 15% of sinus tract cells, sharing the same state mainly with HS lesional epidermal cells.

**Figure 4.**
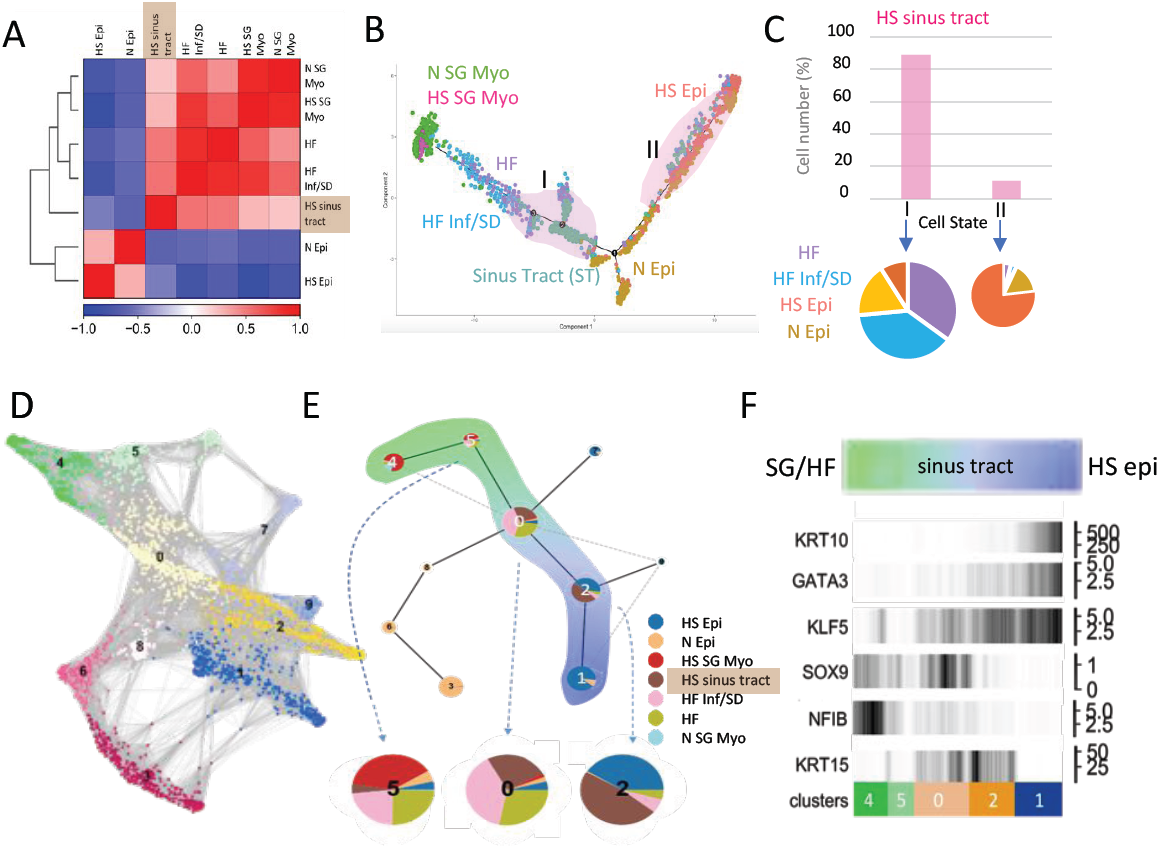
Pseudotime analyses indicate the lineage similarity of HS sinus tracts closest to the infundibulum of apocrine-pilosebaceous unit. (A) Dendrogram, showing correlation matrix between selected epidermis and appendage clusters showed sinus tract cell cluster is grouped with appendage clusters, and closest to HF infundibulum/sweat duct lineage. (B) Pseudotime plot, showing cell trajectories of sweat gland myoepithelial cells (SG Myo), Hair follicle cells (HF), HF infundibulum/sweat duct (HF Inf/SD), sinus tract (ST), and normal (N) and HS surface epidermal (Epi) cells. Pink shaded areas (I and II) indicate the two different cell states that contains HS sinus tract cells. (C) Bar graph, showing the percentage of HS sinus tract cells in cell state I and II. Lower pie charts, showing the percentage of other cell types that shares the same cell state with HS sinus tract cells. (D) PAGA-initialized single cell embeddings, showing the lineage connections between individual cells and 10 Louvain clusters (cell states). (E) PAGA abstracted graph, composed of pie charts for each Louvain cluster, which shows the composition and percentage of different cell types within each cell state. (Lower) Three of the cell states containing HS sinus tract cells are enlarged (5, 0 and 2). Each cell type is color coded as shown on the right. Color-shaded areas, showing the PAGA path containing states 4-5-0-2-1 that is analyzed in (F). (F) PAGA path graph, showing changes in the expression of signature genes defining the fate of appendages and interfollicular epidermis along PAGA path 4-5-0-2-1. Numeric scale on the right, normalized gene expression from Seurat object. Clusters (lower), indicating different cell states along the PAGA paths and the length is in proportion with cell number in the cell state.

Additionally, we use partition-based graph abstraction (PAGA)^28^ to perform lineage analysis for the same clusters and acquired similar results. As shown in Figure 4D, we represented neighborhood graph of single cells Fruchterman-Reingold layout (FR1) (Figure 4D) and then partitioned them into PAGA graph at a coarser resolution (Figure 4E). Connectivity strength between partitions could be further interpreted confidentially and selected to present. Ten partitions (referred as cell states) connect together with three major lineage paths, normal epidermis (3-6-8-), lesional epidermis (1-2-) and appendages (4-5-) with a common intersection at state 0. We further analyzed the composition of cell clusters in each state and found that the majority of the sinus tract cells are distributed in 0 and 2, sharing the lineage similarity with HF and HFinf/SD, and lesional epidermis, respectively. This result is consistent with the results in Figure 4B acquired by Monocle. Further, we selected a path containing state 4-5-0-2-1 for PAGA analysis in single cell resolution and showed a clear transition from skin appendages marker (NFIB and SOX9) high in state 4 gradually transition into state 0, where SOX9 and KLF5 both are present (indication of stem cell fate duality), and then transition into 2 and 1 with increasing levels of interfollicular epidermal markers (KLF5, GATA3, and KRT10), as shown in Figure 6F. Interestingly, KRT15, an epidermal stem cell marker, is present in the state 0 and 2, where the majority of HS sinus tract cells are distributed. Altogether, with two different computational approaches, we conclude that although HS sinus tracts largely exhibit an interfollicular epidermal differentiation program, these cells are in closest lineage relationships with infundibular cells of the pilosebaceous-apocrine unit at the transcriptional level.

**Figure 5.**
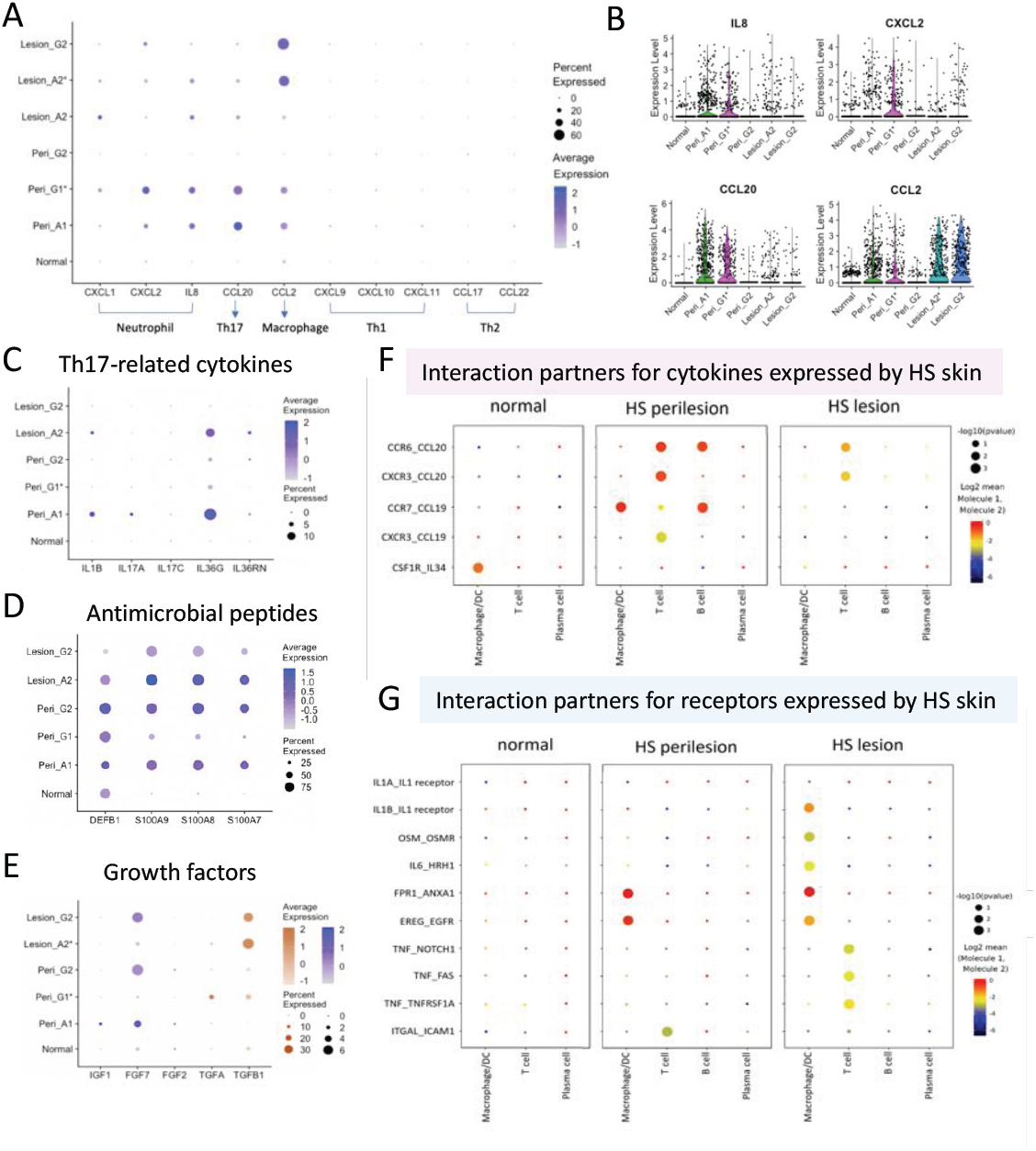
HS perilesional epidermis produces cytokines eliciting Th17 response. (A) Dot plot, showing the percentage of expression and average expression level of key cytokines in the epidermis in different samples. A: axillary skin. G: groin skin. *: basal layer. (B) Violin plots of IL8, CXCL2, CCL20, CCL2, showing their expression levels in the epidermis. (C-E) Dot plot, showing the percentage of expression and average expression level of (C) cytokine genes related to Th17 immune response, (D) antimicrobial peptides, and (E) growth factors. Note that IL36G and IL1B are up-regulated in HS epidermis and variable between patients. FGF7 expression of upregulated in some patients but not all, and TGFB1 is expressed only in the lesional skin. (F-G) Dot plot, showing cytokines-receptor interactions in normal, HS perilesional and lesional skin (Patient A1). Interactions on the left (X_Y), indicating X: gene expressed in Immune cells at the bottom and Y: gene expressed in epidermal cells on the top. (F), showing interactions of cytokines expressed by epidermal cells. (G), showing interactions with receptors expressed by epidermal cells.

**Figure 6.**
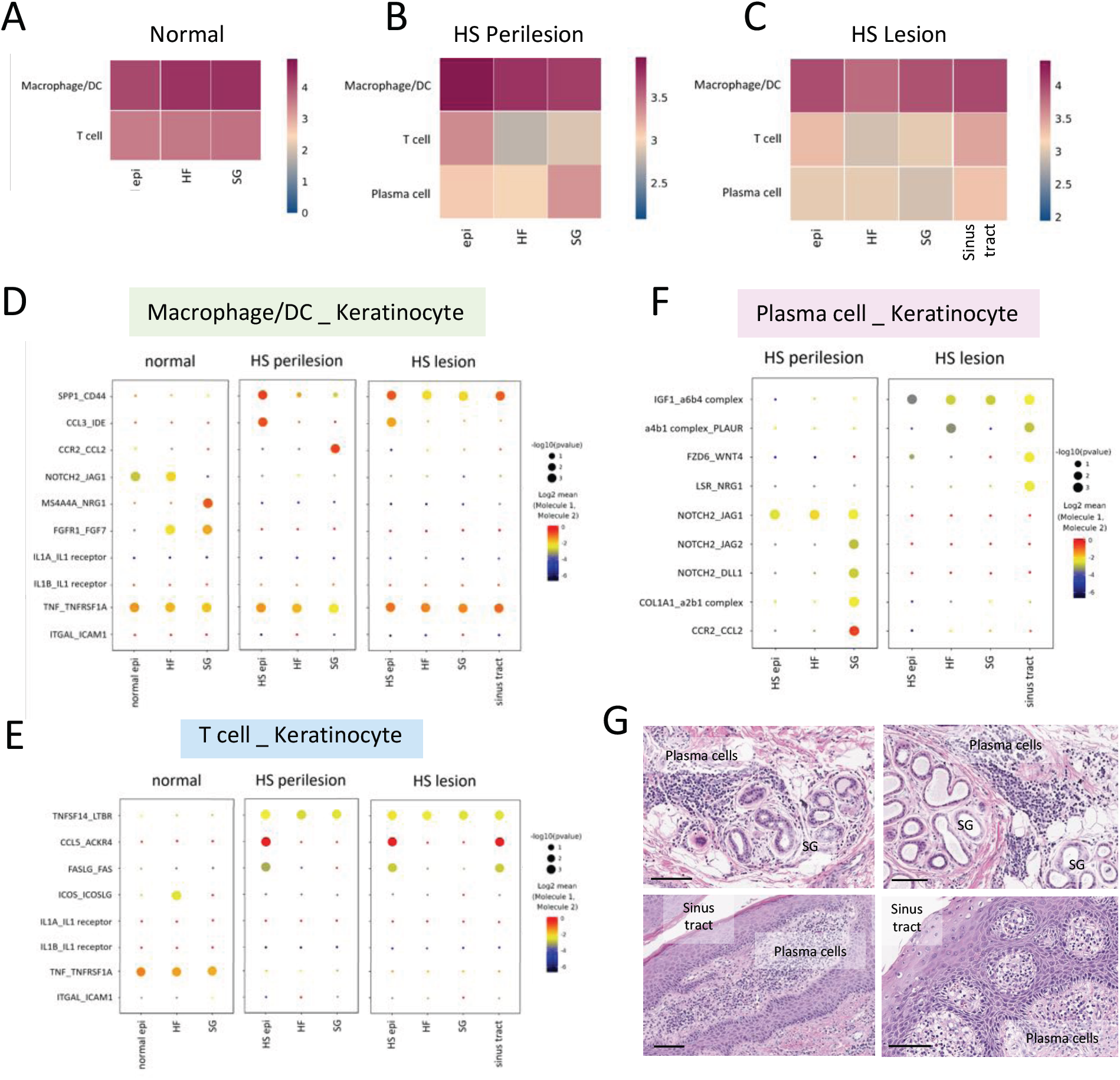
Immunogenicity of skin appendages and sinus tracts in HS pathogenesis. (A-C) Statistical interaction heatmap, showing the intensity of interactions between immune cells and epidermal cells from normal skin (A), HS perilesional skin (B), and HS lesional skin (C). Epi, surface epidermis. HF, hair follicle. SG, sweat gland. DC, dendritic cells. (D-F) Dot plot, showing high-ranked interaction partners between Immune cells, including (D) macrophage/DC, (E) T cell, and (F) plasma cell, with different keratinocyte cell clusters as indicated. Interaction partner labeled on the left (X_Y): X, gene expressed by immune cells; Y, gene expressed by epidermal cells. (G) H&E staining of HS perilesional and lesional skin, showing plasma cells are present in high density near the sweat gland and sinus tract.

### HS perilesional keratinocytes produce cytokines eliciting Th17 response

Using scRNAseq data, we analyzed important cytokines, growth factors, and antimicrobial peptides in normal, perilesion and lesion HS interfollicular epidermis (i.e. excluding clusters from appendages). In Figure 5A, we showed CXCL1, CXCL2, IL8, CCL20 and CCL2 are upregulated in HS epidermis, but not CCL4, CCL5, CXCL9, CXCL10, CCL17, CCL22, confirming that activated keratinocytes in HS skin mediate recruitment of T cells associated with IL17 responses, as well as macrophages and neutrophils, whose important roles had been implicated in HS^2,29,30^. Importantly, we detected higher levels of IL8, CXCL2, and CCL20 in perilesional epidermis than in lesional epidermis (Figure 5B), suggesting that these are early responders for epidermal abnormality, and likely downregulated when severe inflammation has been established in lesional skin. With regards to IL17 response-related chemokines, we detected low levels of IL1B, IL36G, IL36RN, and surprisingly IL17A in patient Ax1 (Figure 5C), whose CCL20 level is also particularly high. We also detected expression of antimicrobial peptides including DEFB1, S100A7, S100A8, S100A9, which also show significant expression already in perilesional skin (Figure 5D). With regards to growth factors, FGF7 and TGFB1 are detected in HS epidermis. In particular, there are much higher levels of TGFB1 in lesional skin than in peri-lesional skin, indicating that TGFB1 is a late responder in HS and correlates with the disease stage when fibrotic scarring occurs extensively (Figure 5E).

Next, we used CellPhoneDB v2.0^31^ to investigate whether and which immune cells express the receptors for the cytokines expressed by epidermal cells, and to analyze the potential interactions between keratinocytes and immune cells by permutations with statistical confidence. We used patient Ax1 (whose epidermis expressed the highest level of CCL20) as an example and analyzed the potential interactions with immune cells present in the same sample. As shown in Figure 5F, we detected high levels of CCL20 interaction with CCR6 and CXCR3 expressed by T cells in the perilesional skin. This interaction is lower in lesional skin, mainly due to lower expression of CCL20 in the lesional epidermis (Figure 5A). CCL19 and CCL20 are expressed at a higher level in perilesional skin and are both important for lymphocyte chemoattraction. In addition, IL34 is expressed in the normal epidermis and not HS lesional epidermis, suggesting that it may mediate monocyte/macrophage survival during homeostasis and that this interaction may be lost in HS skin. Further, we sought to understand how epidermal cells may respond to cytokines produced by immune cells by analyzing the receptors/interaction partners that are expressed by epidermal cells. As shown in Figure 5G, epidermal cells in the perilesional skin express ANXA1 and EGFR, which could have strong interactions with FPR1 and EREG, expressed by macrophage/DC, respectively. These interactions continue in the more severe stages (lesional skin). In lesional skin, the epidermal cells also express IL1R, OSMR, and HRH1, that could receive the IL1B, OSM, IL6 produced by macrophages/DCs, which are critical for augmentation of inflammatory response. With regards to interactions with T cells, HS lesional epidermal cells express TNFRSF1A and ICAM1, that could interact with TNF and ITGAL (LTA-1 or CD11a), respectively. It is particularly interesting that ITGAL_ICAM1 interaction occurs only in perilesional skin, while TNF_TNFRSF1A (and other interaction partners FAS and NOTCH1) interactions occur in lesional skin, which may have important implications on therapeutic strategies.

### Immunogenicity of skin appendages and sinus tracts in HS pathogenesis

In order to delineate the roles of different skin appendages and sinus tracts in HS pathogenesis, we used CellPhoneDB v2.0 to further analyze potential immune-epithelial communication networks in normal, HS perilesional and lesional skin (from patient Gr2), with a specific focus on the comparisons between interfollicular epidermis (epi), hair follicle (HF), sweat gland (SG) and sinus tract (ST). In Figure 6, we showed in the heatmaps the interaction intensity of possible immune-epidermal combinations in normal (Figure 6A), HS perilesional (Figure 6B) and lesional (Figure 6C) skins. We found that macrophage/DCs have strongest interactions with all epidermal clusters in all three stages. The potential interactions are shown in Figure 6D, indicating that they use different sets of ligand-receptor to interact with epidermal cells in normal vs HS skin. In particular, CCL3_IDE and CCR2_CCL2 interactions may occur between macrophage/DCs with interfollicular epidermis and sweat gland, respectively, and both of which are important in the establishment of chemotactic gradients. With regards to T cells, they showed high levels of interactions with HS perilesional epidermis compared with appendages (HF and SG); however, in the lesional skin, T cells have even higher levels of interaction with keratinocytes in the sinus tract than in the epidermis. This may result more T cells being recruited to the sinus tract, which is consistent with our quantification of immune cells in various regions of HS lesional skin shown in Sup Figure 1. As shown in Figure 6E, amongst these potential interactions, TNFSF14_LTBR could be a general interaction that occur in HS, while CCL5_ACKR4 and FASLG_FAS are more specific to interfollicular epidermis and the sinus tract. Interestingly, we found in this patient (Gr2) that TNF_TNFRSF1A interaction occurs between macrophage and epidermis, and not between T cells and epidermis as in patient Ax1 (Figure 5G), suggesting that fundamentally different immune responses may occur in different HS patients.

Further, we investigated the presence of plasma cells in the HS skin and their potential interactions with different types of keratinocytes, which has been largely neglected in the literature. Intriguingly, we noticed a high level of interaction between plasma cells and sweat glands in HS perilesional skin (Figure 6B) as well as interaction between plasma cells and sinus tracts in the HS lesional skin (Figure 6C). As shown in Figure 6F, we found that sweat gland cells express JAG1, JAG2, DLL1, that could be interacting with NOTCH2 expressed by the plasma cells in the perilesional skin. In addition, SG cells express high level of CCL2, which could serve as a strong chemoattractant for CCR2-expressing plasma cells. As for the lesional skin, keratinocytes in the sinus tracts express higher levels of WNT4 and NRG1, which could contribute to the interaction with FZD6 and LSR expressed by plasma cells, respectively. In Figure 6G, we showed that plasma cells are recruited to the vicinity of sweat glands and sinus tract, supporting the notion that sweat gland and sinus tract may have unique roles in recruitment of plasma cells in HS skin.

## Discussion

Previous work based on histological observations concluded that Hidradenitis Suppurativa is a disease derived from follicular occlusion^19–21^ and it has also been proposed that the unique features in immune and barrier functions make apocrine sweat gland-bearing skin regions prone to HS development^32^. Here we finely dissect the roles of different types of keratinocytes of the pilosebaceous-apocrine unit in the pathogenesis of HS. First, we showed that keratinocytes lining HS sinus tracts express high levels of genes specific for sweat and sebaceous glands, albeit their morphology and differentiation program resembles interfollicular epidermis. Secondly, we showed that stem cell infidelity/duality exists significantly in the HS sinus tracts, similar to neoplastic and wound conditions^17^, which contributes to the epidermal abnormality in HS pathogenesis. Using single cell RNAseq analyses, we identified changes in gene expression in different types of keratinocytes during HS progression, by comparing normal, perilesional and lesional skin, and we showed that cells lining HS sinus tracts share the closest lineage relationship with infundibulum of the pilosebaceous-apocrine unit. Altogether, our data support the model that epidermal stem cells residing in the infundibulum exhibit cell fate duality, and that loss of appendage fate (SOX9) may trigger epigenetic changes^16,33^ that lead to the formation of sinus tracts with features resembling interfollicular epidermis while still maintaining the potential to express glandular signatures.

It has been demonstrated in mouse models that the loss of key transcription factor Foxc1 in HF stem cells leads the inability to maintain their cells fate^34,35^, and further, the reduction of E-cadherin^35^. Moreover, in the case of complete loss of E-cad in HF stem cell niche, Cldn1 expression is down-regulated, while Ccl2 expression is upregulated, which leads to immune cell recruitment, independent of bacterial invasion^36^. These changes are consistent with the potential development of the sinus tract in HS: FOXC1 and CDH1 expression is downregulated, and CCL2 is upregulated in the HS sinus tract compared with normal infundibulum and HS infundibulum (Figure 3E and Sup Figure 8). Therefore, the stem cell fate perturbation in the HS sinus tract likely contributes to the initiation of the immune response that continues to exacerbate. This may also explain the higher level of immunogenicity in the HS sinus tracts as we demonstrated in Sup Fig 1 and Figure 6 C-F, compared to interfollicular epidermis, which makes HS more difficult to treat.

With regards to the role of hair follicle (HF) and sweat gland (SG) stem cells in HS pathogenesis, our data demonstrated that the direct conversion into sinus tracts is unlikely and that similar changes occur in both appendages in the HS lesions. First, HF and SG in the lesional skin do not appear to have as much extensive proliferation as in the sinus tracts. Proliferation has only been observed in infundibulum and sweat duct (sup Figure 9). Second, in pseudotime lineage analyses, lesional SG cells exhibit a relationship closer to infundibulum/sweat duct cells that largely overlaps with sinus tracts, which may be an indication of gradual loss of specificity (sup Figure 10). Third, both HF and SG in the HS lesion exhibit decreased levels of stem cell marker and AP1 family members (Figure 3G-H and sup Figure 7), and increased level of IFN response genes (sup Figure 11), consistent with recent work^37^. In addition, our data suggests intriguing and unique roles of sweat glands in HS pathogenesis: 1) we detected upregulation of ribosomal proteins in the lesional SG, including RPL7 (Figure 3G), which was shown to be an autoantigen in lupus^38^; 2) SG cells in perilesional skin display an unexpected high level of interaction with plasma cells, in particular, using JAG1, JAG2, DDL1 to interact with NOTCH2 expressed by plasma cells, as well as expressing high level of CCL2 to interact with CCR2 expressed by plasma cells (Figure 6F). These interactions may likely contribute to the specific recruitment of plasma cells to the vicinity of sweat glands as shown in Figure 6G. Our data suggests that although sweat glands do not directly contribute to sinus tract formation, they may have the potential to elicit an autoimmune response.

Current ongoing HS clinical trials are designed to repurpose existing monoclonal therapeutics to block IL17A-(Secukinumab, Bimikizumab, Ixekizumab), TNFα-(Adalimumab, Etanercept, Infliximab), IL1α-(Anakinra and Bermekimab), and LFA1-(Efalizumab) mediated inflammation^39^. Since HS pathogenesis had not been thoroughly characterized, our work thus provides important implications with regards to efficacy of these treatments. First, we detected the expression of CCL20, and surprisingly low level of IL17A, in HS keratinocytes, but only in one patient (Ax1). In addition, we showed that the critical interactions to initiate Th17 response, CCL20_CXCR3 and CCL20_CCR6, occur at higher levels in perilesional skin than in lesional skin, which may be attributable to downregulation of CCL20 in lesional epidermis (Figure 5 A and F). This same patient also has the highest number of IL17A+ T cells (data not shown). Together, these results suggest a spectrum of variable levels of IL17 response could be elicited amongst HS patients. Secondly, we showed that the level of TNFα-IL1β axis response in HS epidermis is higher in lesional than perilesional skin (Figure 5G and 6D), which supports the current role of TNFα-IL1β blockade in the clinical management of disease. Third, we showed that IL1A_IL1R interactions are not present in our HS samples; however, IL1B_IL1R is detected in Ax1 patient, whose epidermis also express low level of IL1B, which indicates that blocking IL1R may be effective in some patients, while blocking IL1A may not. Again, this is corroborated by the contrasting results of small scale clinical trials of IL-1B and IL-1a blockade in clinical disease^40–42^. Lastly, we showed that ITGAL(LFA-1)_ICAM1 interaction is minimally detectable in perilesional epidermis of one patient (Ax1), and not others, confirming that LFA1 blocking may not be effective for HS treatment^43^.

Altogether, our work using Hidradenitis Suppurativa as a human inflammatory skin disease model illustrates the crucial roles of stem cell fate maintenance in pathological epidermal remodeling, as well as the contribution of different compartments of pilosebaceous-apocrine units to its unique and complex pathological responses.

## Materials and Methods

### Antibodies

Chicken anti-keratin 14 Biolegend Cat#906001

Rabbit anti-SCD1 Abcam Cat#ab39969

Rabbit anti-keratin 77 Abcam Cat#ab106847

Mouse anti-CEA Invitrogen Cat#MA5-15070

Mouse anti-Sox10 R&D Systems Cat#MAB2864

Rabbit anti-Sox9 Millipore Cat#AB5535

Goat anti-Klf5 R&D Systems Cat#AF3758

Rabbit anti-Cldn1 ThermoFisher Cat#71-7800

Rabbit anti-Timp1 Abcam Cat#Ab211926

Rabbit anti-Ki67 Abcam Cat#Ab15580

Rabbit anti-Lhx2 (kindly provided by Fuchs lab, Rockefeller University)

Rat anti-CD49f BD Pharminogen Cat# 55734

Rabbit anti-Keratin 10 Biolegend Cat#905404

Donkey anti-rabbit Alexa Fluor 488 Jackson ImmunoResearch Cat#711-545-152

Donkey anti-chicken Alexa Fluor 647 Jackson ImmunoResearch Cat#703-605-155

Donkey anti-Rat Alexa Fluor 647 Jackson ImmunoResearch Cat#712-605-150

Donkey anti-goat Cyanine Cy3 Jackson ImmunoResearch Cat#705-165-147

Donkey anti-mouse Rhodamine Red-X Jackson ImmunoResearch Cat#715-295-150

Donkey anti-chicken Alexa Fluor 488 Jackson ImmunoResearch Cat#703-545-155

Donkey anti-rabbit Cyanine Cy3 Jackson ImmunoResearch Cat#711-165-152

DAPI solution Fisher Scientific Cat#D1306

### Chemicals

HBSS Fisher Scientific Cat# MT21021CV

Collagenase Type I-A Sigma Aldrich Cat# C2674

Hyaluronidase Sigma Aldrich Cat#H3506

Trypsin/EDTA Corning Cat# MT25053CI

ACK Lysing Buffer Fisher Scientific Cat#A1049201

Fetal Bovine Serum Fisher Scientific Cat# 35-010-CV

### Software and Algorithms

R R Development Core Team, 2008 http://www.R-project.org

Seurat CRAN https://satijalab.org/seurat/

Monocle Bioconductor http://cole-trapnell-lab.github.io/monocle-release/docs/

iCellR CRAN https://github.com/rezakj/iCellR

PAGA (partition-based graph abastraction) CRAN https://github.com/theislab/paga

CellPhoneDB v2.0 https://www.cellphonedb.org/

### Pipeline

**link to GitHub**

### Experimental Model and Subject Details

#### Human Specimens

Patient samples were collected from severe HS cases screened by the Hansjorg Wyss Department of Plastic Surgery. Protocols for sample collection were reviewed by the Institutional Review Board (IRB) at NYU Langone (s19-00841). Inflamed regions, characterized by nodules, abscesses, and draining fistula, were surgically excised from axillary and groin regions. Following the collection of samples, patient tissue was dissected to identify epidermal and dermal regions, hair follicles, and sinus tracts.

### Method Details

#### Preparation of patient samples for tissue sectioning and immunofluorescent staining

Upon receipt from surgery, patient tissue samples were dissected to isolate regions of interest, such as fistulae, sinus tracts, and inflamed epidermis. The dissected area was freshly embedded in OCT compound (Sakura Finetek USA, Inc., Torrance, CA) and immediately frozen on dry ice. OCT blocks were kept at −80°C until needed for sectioning. OCT blocks were cryosectioned at 10 μm thickness using the Leica cryostat and underwent immunofluorescent staining. Slides that were not immediately used for staining were kept at −20°C until needed.

#### Immunofluorescent staining and imaging

Thawed cryo-sectioned slides were briefly fixed with 4% PFA in PBS, then washed three times in 1X PBS. Sections were then permeabilized in 0.1% PBST (Triton X-100 (Fisher Scientific, Fair Lawn, NJ), and 1X PBS) before blocking for one hour with blocking buffer (Bovine Serum Albumin (Fisher Scientific, Fair Lawn, NJ), Normal Donkey Serum (Jackson ImmunoResearch Inc., West Grove, PA), Normal Goat Serum (Jackson ImmunoResearch Inc., West Grove, PA), Triton X-100 (Fisher Scientific, Fair Lawn, NG), and 1X PBS). Slides were incubated with the following primary antibodies diluted in blocking buffer overnight at 4°C – chicken anti-keratin 14 (1:800, Biolegend 906001), rabbit anti-SCD1 (1:500, Abcam, ab39969), rabbit anti-keratin 77 (1:500, Abcam, ab106847), mouse anti-CEA (1:100, Invitrogen, MA5-15070), mouse anti-Sox10 (1:800, R&D Systems, MAB2864), rabbit anti-Sox9 (1:1000, Millipore, AB5535), goat anti-Klf5 (1:500, R&D Systems, AF3758), rabbit anti-Cldn1 (1:100, ThermoFisher, 71-7800), rabbit anti-Timp1 (1:500, Abcam, ab211926), rabbit anti-Ki67 (1:500, Abcam, ab15580), rabbit anti-Lhx2 (1:1000, Fuchs lab), rabbit anti-NFIX (1:1000, Fuchs), rabbit anti-NFIB (1:1000, Fuchs), rat anti-CD49f (1:200, BD Pharminogen, 55734), rabbit anti-Keratin 10 (1:100, Biolegend, 905404). The following day, the slides were washed three times with 1X PBS before incubating the slides in the following secondary antibody for one hour at room temperature, including the addition of DAPI – donkey anti-rabbit Alexa Fluor 488 (1:800, Jackson ImmunoResearch, 711-545-152), donkey anti-chicken Alexa Fluor 647 (1:800, Jackson ImmunoResearch, 703-605-155), donkey anti-Rat Alexa Fluor 647 (1:800, Jackson ImmunoResearch, 712-605-150), donkey anti-goat Cyanine Cy3 (1:200, Jackson ImmunoResearch, 705-165-147), donkey anti-mouse Rhodamine Red-X (1:200, Jackson ImmunoResearch, 715-295-150), donkey anti-chicken Alexa Fluor 488 (1:800, Jackson ImmunoResearch, 703-545-155), donkey anti-rabbit Cyanine Cy3 (1:200, Jackson ImmunoResearch, 711-165-152). The slides were washed in PBS a final time before applying coverslips with ProLong Gold antifade reagent (Invitrogen by ThermoFisher Scientific, Eugene, OR). IF stained cryo-sectioned slides were then imaged using the Zeiss Axio Observer at 20x objective lens and were processed using Fiji.

#### Preparation of epidermal single cell suspensions

Samples from four different patients were collected, including perilesion and lesional areas, as well as two samples from patients without HS. Entire excisions were collected from human patients and placed directly into sterile Dulbecco’s Phosphate Buffered Saline (DPBS, 1X w/o Ca2+ Mg+ (Corning, Manassass, VA)) on ice within an hour of the surgery. Areas of interest, focusing on sinus tracts and inflamed epidermis, as well as perilesional areas, were dissected out of the larger sample to 1cm biopsies. These biopsies were then dissociated into a single cells suspension using an enzyme mix of Hanks’ Balanced Salt Solution (HBSS, 1X w/o Ca2+ Mg+ (Corning, Manassass, VA)), Collagenase at 20 mg/ml (Sigma Aldrich, St. Louis, MO), and Hyaluronidase at 20mg/ml (Sigma Aldrich, St. Louis, MO). The samples were incubated at 37°C on a shaking plate at 100 rpm. After 1 hour, the samples were agitated with dissecting scissors to further the digestion process. The samples incubated for 30 more minutes, of which the final 10 minutes Trypsin/EDTA (Corning, Manassass, VA) was added to finalize the digestion process. The suspension was then washed in 4% FBS/DPBS (Corning, Manassass, VA) and filtered through a 40 μm cell strainer and spun down at 330 g at 4°C for 10 minutes. Red blood cell lysis using ACK Lysis Buffer (Gibco, Life Technologies, Grand Island, NY) was then performed, then cells were washed and spun as before.

#### 10x Single Cell RNA sequencing

Single cell suspensions of approximately 1.0e6/ml in PBS were submitted to the Genome Technology core facility at NYU Langone. The following supplies were used at the manufacturer’s instructions: Chromium Single cell 3’ Library and Gel Bead Kit v2 Chromium Single Cell 3’ Library & Gel Bead Kit v2 (PN-120237), Chromium Single Cell 3’ Chip kit v2 (PN-120236_ and Chromium i7 Multiplex Kit (PN-120262). Once the cDNA libraries were generated, they were subject to HiSeq 2500 (illumina) sequencing.

#### scRNA-seq data analysis

The live cells from a total of nine different patient samples, including 6,947 cells from normal axilla (n=1), 1,133 and 7,668 cells from HS axilla lesion replicates (n = 2), 1,561 cells from HS axilla perilesion (n = 1), 3,993 cells from normal groin (n = 1), 2,776 and 7,386 cells from HS groin lesion replicates (n = 2), and 2,010 and 5,047 cells from HS groin perilesion replicates (n = 2) had their transcriptomes sent for sequencing. After confirming the integrity of the cDNA, quality of the libraries, number of cells sequenced and mean number of reads per cell, as a quality control, we used the cellranger package to map the reads and generate gene-cell matrices. A quality control was then performed on the cells to calculate the number of genes, UMIs and the proportion of mitochondrial genes for each cell using Seurat v3.0 (Butler et al., Nature Biotechnology 2018: Stuart, Butler, et al., Cell 2019) and the cells with low number of covered genes (gene-count < 500) and high mitochondrial counts (mt-genes > 0.1 or 10%) were filtered out. The data was then normalized (log normalized), scaled, and variable features (dispersed genes) were used to perform principle component analysis (PCA). The first 40 PCs were used to find neighbors and clusters, using the FindNeighbors and FindClusters functions (resolution at 0.5). We then performed Stochastic Neighbor Embedding (t-SNE) and Uniform Manifold Approximation and Projection (UMAP) on the first 40 PCs. Marker genes per clusters were found. Cell trajectories were constructed from selected epidermal and appendage clusters using Monocle v2.0 (Trapnel, Cacchiarelli et al, Nature Biotechnology, 2014). The data went under unsupervised clustering, then determined the ordering genes based on differential gene test. From there, the cell trajectory could be plotted to show the lineage of epidermal, appendage, and sinus tract cell types. Cell trajectories were also constructed by another tool, partition-based graph abstraction (PAGA) (Theis lab, Alexander Wolf et al, Genome Biology, 2019), using the same selected clusters. Parameters including n_neighbors = 80 and resolution =0.7 were determined to generate abstract of cell trajectories. The package iCellR (v1.3, https://github.com/rezakj/iCellR) was used to generate interactive plots trajectory plots in order to compare gene expression of different cell types in different cell states.

We extracted patient-specific Seurat project from keratinocytes and immune cells and further re-cluster and annotate by Seurat. We then input these individual extracted raw data into online publicly available repository (CellPhoneDB v2.0), seeking for curated receptors, ligands and their interactions.

## Data availability

Single cell RNA sequencing data used to support the findings of this study are in the process to be deposited in Gene Expression Omnibus (https://www.ncbi.nlm.nih.gov/geo/). All data acquired and generated by the lab are recorded in the manuscript and the supplementary materials, or available from the corresponding author upon reasonable request. The code/Rmarkdown files for the analyses reported in this paper will be available at (GitHub link).

## Acknowledgements

We are grateful to the kind support and comments from Drs. Rodriguez E, Ceradini D, Rabbani P, Ito M and Cadwell K at NYU Langone, and insightful clinical advices from Drs. Krueger J. and Frew J. We would also like to thank Dr. Kasper M for critical aid in single cell analyses, as well as Krylyuk H, Maharaj A, Dr. Markopoulos C for preliminary data processing. We thank facilities at NYU Langone: Experimental Pathology Research Laboratory (S10 OD021747), the Applied Bioinformatics Laboratories, and the Genome Technology Center (P30CA016087). C.L. and lab members are supported by start-up fund from Hansjörg Wyss Department of Plastic Surgery. W.Y. was supported by Yen Tjing Ling Medical Foundation.

## Author contributions

C.L. conceived the project and designed experiments. M.M., M.L., W.Y. and C.M.M. performed experiments and analyses. J.R. assisted tissue acquisition and manuscript writing. A.K-J. provided advanced computational analyses. E.C. performed surgery and patient care. C.L. wrote the manuscript.

