## Supplementary Figures for "Defining Epidermal Stem Cell Fate Infidelity and Immunogenicity in Hidradenitis Suppurativa at the Single-Cell Resolution"

**Sup Figure 1.**

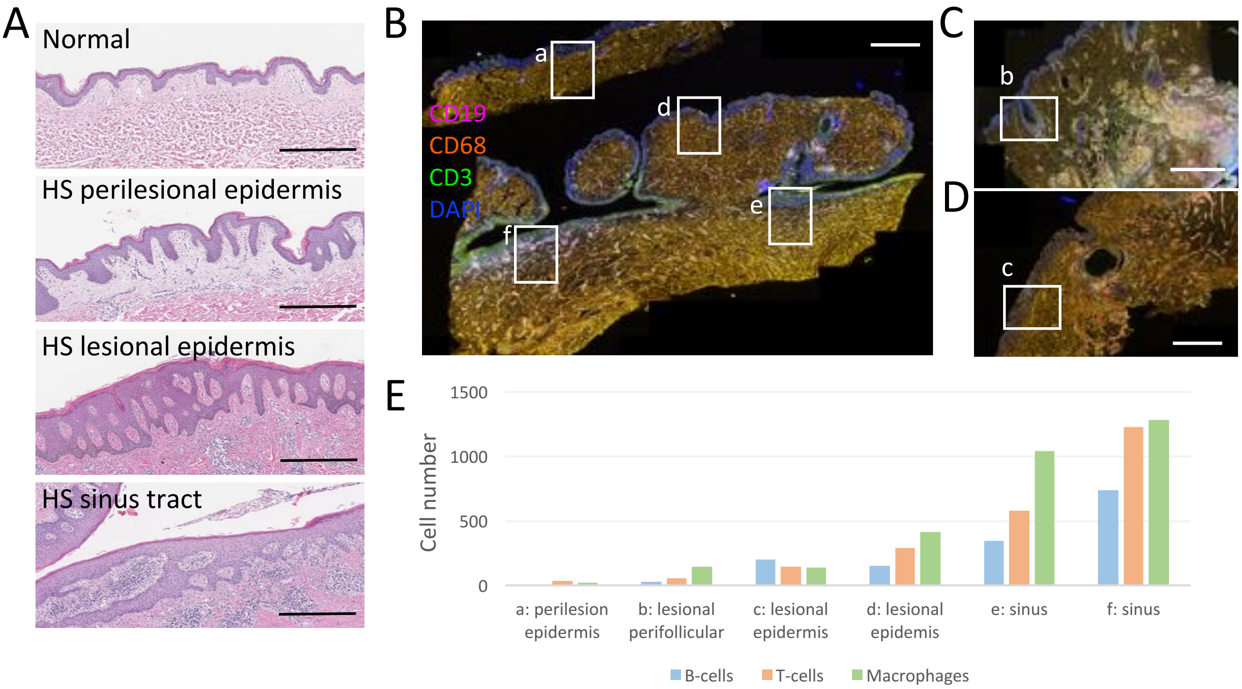

**Sup Figure 1. Epidermal abnormality occurs prior to immune cell recruitment.**
(A) From top to bottom, H&E staining of human normal skin, HS perilesional, lesional and sinus tract epithelium. Scale bars, 500um. (B-D) Representative pictures of CD3, CD19, CD68, and DAPI staining in multiplexed images. White boxes annotate different areas of HS lesional and perilesional skin. a, HS perilesional epidermis. b, lesional perifollicular area. c-d, lesional epidermis. e-f, lesional sinus tract. Scale bars, 2mm.(E) Quantification of inflammatory cells in different boxed areas in (B-D). Noted that perilesional epidermis has very little immune cell infiltration while exhibiting significant hyperplasia, and that in HS lesional skin, sinus tract areas contains significant higher number of immune cells than surface epidermal areas.

### Sup Figure 2.

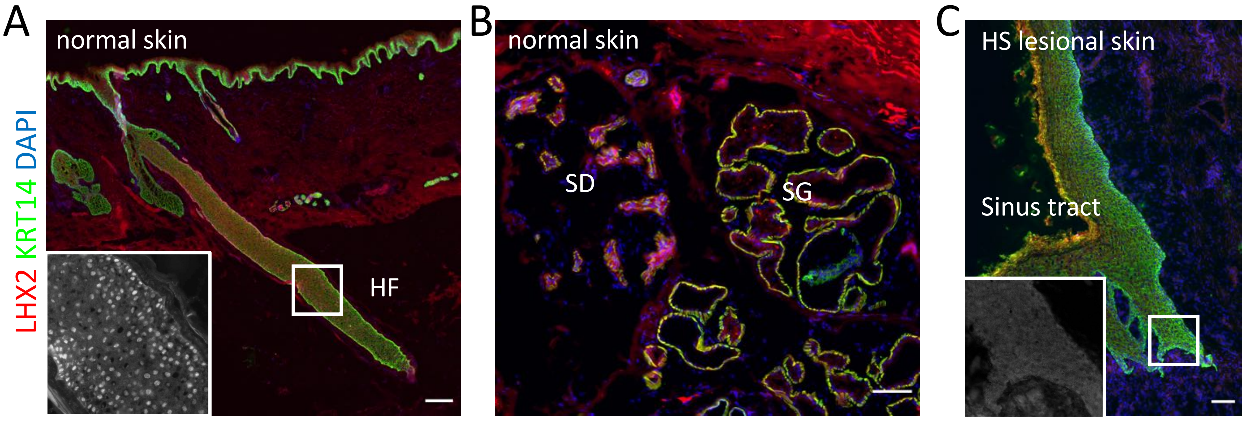

**Sup Figure 2. Immunofluorescent staining for LHX2 in (A-B) human normal skin and (C) HS lesional skin showing sinus tract.**
Noted in normal skin, LHX2 is specifically expressed in hair follicle, but not in sweat gland; and in HS sinus tract, LHX2 expression is not detectable. Scale bar, 200 µm in (A) and 100 µm in (B-C). HF, hair follicle. SD, sweat duct. SG, sweat gland. Box in (A), high magnification of hair follicle, showing nuclear expression of LHX2. Box in (C), high magnification for protrusion front of sinus tract, showing no LHX2 expression.

**Sup Figure 3.**

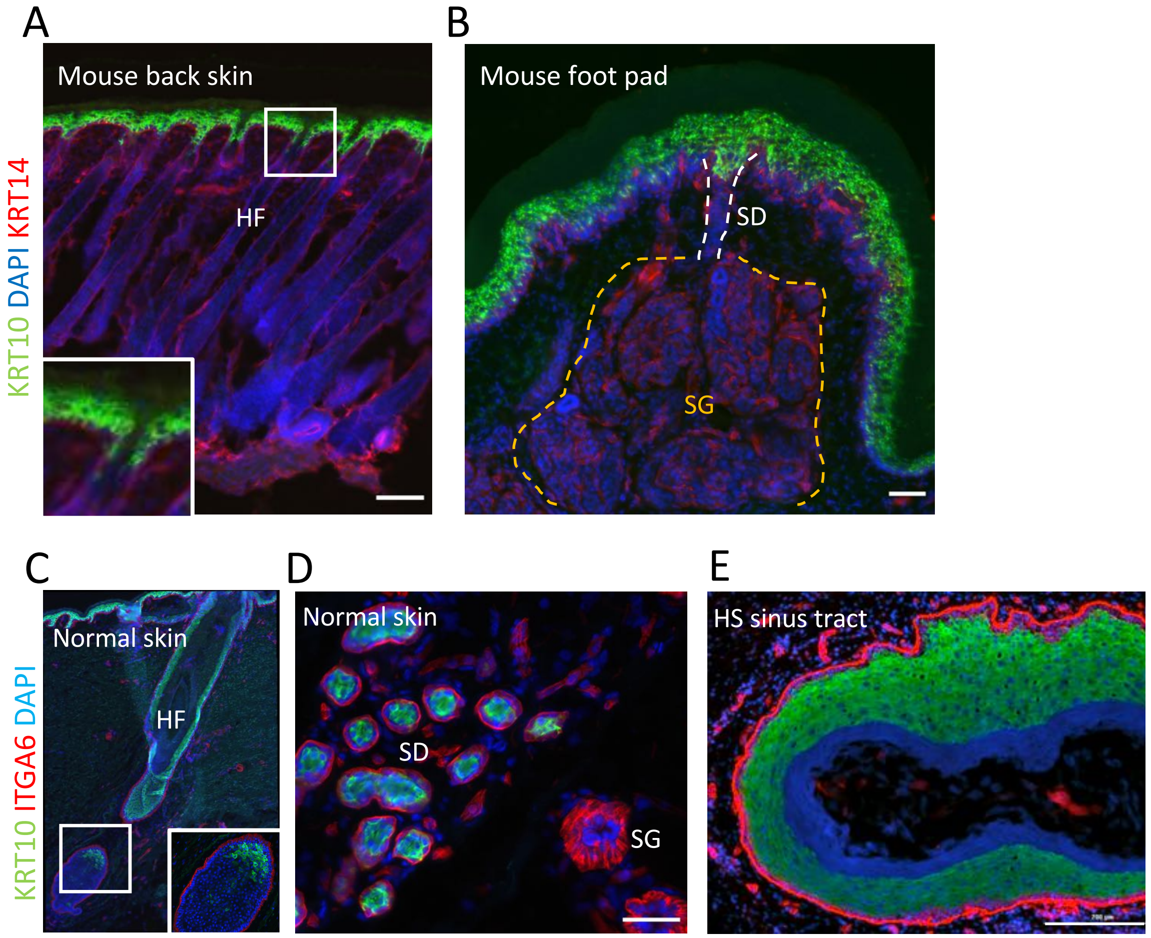

**Sup Figure 3. Immunofluorescent staining for KRT10 in (A) mouse back skin, (B) mouse foot skin, (C-D) human normal skin, and (E) HS sinus tract.**
Noted in mouse skin, where KRT10 expression is restricted to the surface epidermis, and only the very top part of appendages: hair follicle (HF) infundibulum (A) and sweat ducts (B). However, in human skin, KRT10 expression extend much deeper in HF (C) and the entire sweat duct, suggesting that human skin appendages harbor more cells that present dual fate identity (ie. appendage and interfollicular epidemis). Scale bar in A, 100um; B, 50um; D, 50um; E, 200um. Box in (A), higher magnification of upper part of infundibulum. Dashed line in (B), outlining the areas for sweat duct (SD) and sweat gland(SG), respectively. Box in (C), higher magnification for lower part of human HF with cells still express high level of KRT10.

**Sup Figure 4.**

A

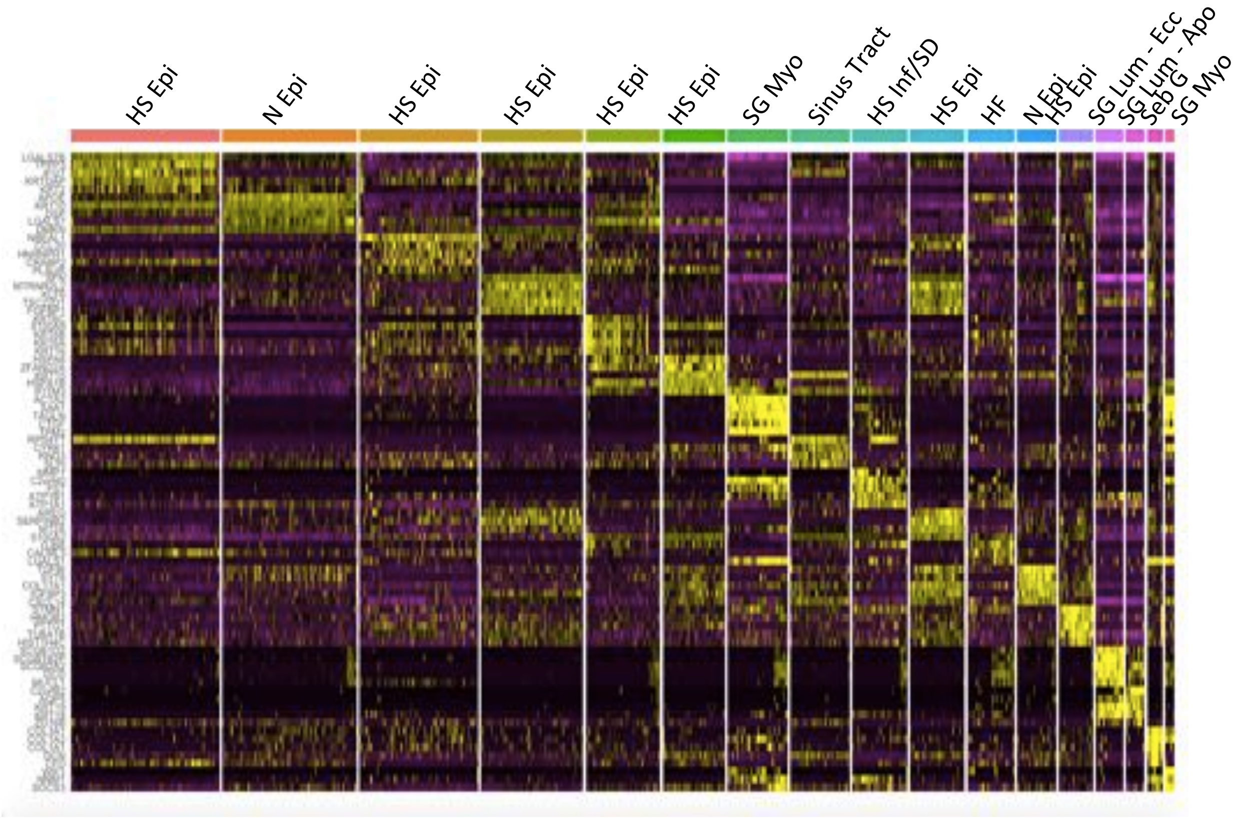

B

| HS Epidermis | Normal Epidermis | HS Epidermis | HS Epidermis | HS Epidermis | HS Epidermis | SG - Myo | HS Sinus Tract | HF Inf/ SD |
|---|---|---|---|---|---|---|---|---|
| LGALS7B | APOE | RPL36A | FOSL1 | SERPINB4 | SLC47A2 | ACTA2 | RPS4Y1 | AC006262.5 |
| RPS4Y1 | KRT10 | AREGB | SFN | SERPINB3 | RAMP1 | SAA1 | KRT15 | MMP7 |
| TPPP3 | PERP | RPS2 | MTRNR2L8 | KRT6C | LMO1 | TAGLN | WNT3 | KRT23 |
| KRT10 | DSP | GLTSCR2 | RPS26 | RHOV | HSPA6 | CTGF | IFITM1 | C2orf40 |
| LY6D | LGALS7 | NBEAL1 | MTRNR2L2 | LGALS7 | SERPINH1 | TPM2 | DLK2 | FST |
| KRTDAP | SCEL | RPS28 | KLF6 | HSPA1B | WNT7A | MYLK | THBS2 | PNLIPRP3 |
| KRT1 | RPS4X | RPS27 | STX11 | IGLL5 | IFITM1 | MYL9 | GSTT1 | FGF7 |
| IFI27 | YBX3 | ITM2B | TSC22D1 | HSPA6 | ZFAND2A | SAA2 | KRT31 | MYBPC1 |
| DBI | MT-ND2 | YBX1 | DDX21 | HSPA1A | KRT15 | ACTG2 | TIMP1 | MFAP5 |
| DMKN | DMKN | RPL39 | ZNF57 | ZGAND2A | ASS1 | FBXO32 | PLAT | MIA |

| HS Epidermis | HF | Normal Epidermis | HS Epidermis | SG Lum - Ecc | SG Lum - Apo | Seb G | SG - Myo |
|---|---|---|---|---|---|---|---|
| POSTN | LPHN3 | TGFBI | KIAA0101 | SCGB1D2 | C2orf82 | COL6A3 | FAM107A |
| AREG | FAM132A | ASS1 | TK1 | SCGB1B2P | S100A1 | TMEM176B | WIF1 |
| SLCO4A1 | LHX2 | SYT8 | UBE2C | PIP | TESC | OLFML3 | PRKD1 |
| PANX1 | FXYD6 | COL17A1 | CDK1 | AZGP1 | TMEM213 | OLFML2B | PCP4 |
| TNFRSF12A | PTN | IGFBP3 | TOP2A | KRT19 | CLDN10 | TWIST1 | SYNM |
| SERPINB2 | CYP1B1 | KRT14 | BIRC5 | SCGB2A1 | PPP1R1B | OLFML1 | CNN1 |
| ERRFI1 | CARD18 | GDPD2 | CDKN3 | TSPAN8 | NCALD | SLIT3 | STGALNAC5 |
| FOSL1 | ABI3BP | FAM134B | RRM2 | LRRC26 | AQP5 | PRRX1 | SLC2A4 |
| PHLDB2 | APOC1 | KRT5 | MKI67 | OBP2B | CA6 | TNXB | ACTG2 |
| ARID5B | CREB5 | CXCL14 | PBK | CLDN3 | ROPN1B | F10 | PPP1R14A |

**Sup Figure 4.**

C

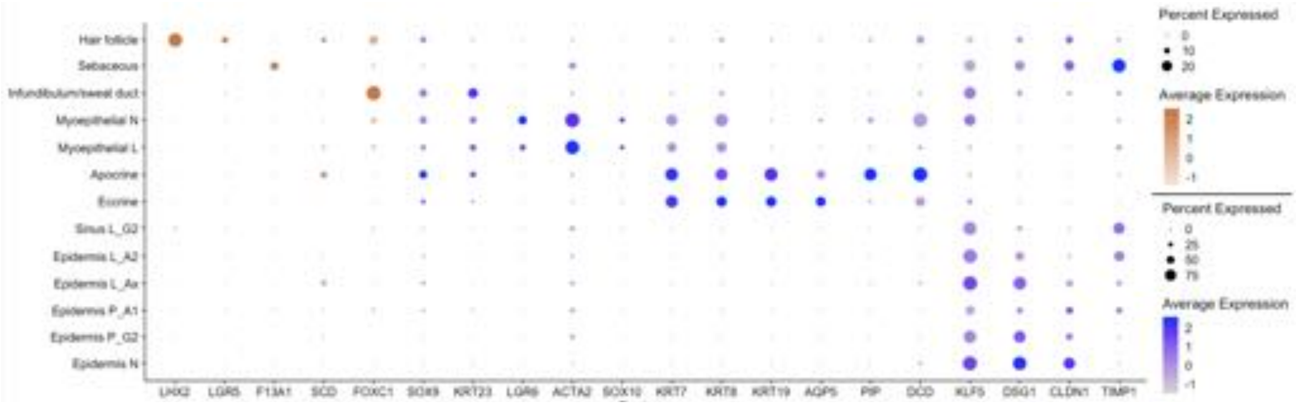

**Sup Figure 4. single cell RNAseq analyses for all the keratinocyte clusters.**
(A) Heatmap of all of the keratinocyte clusters. (B) List of the top genes in each clusters. (C) Dot plot of known markers for various keratinocyte cell types (from mouse models) facilitate the identification of cluster identify in human skin.

**Sup Figure 5.**

A

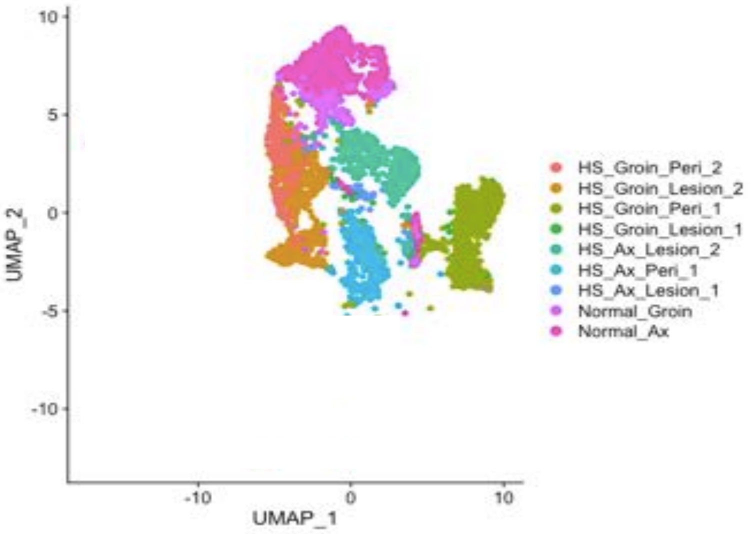

B

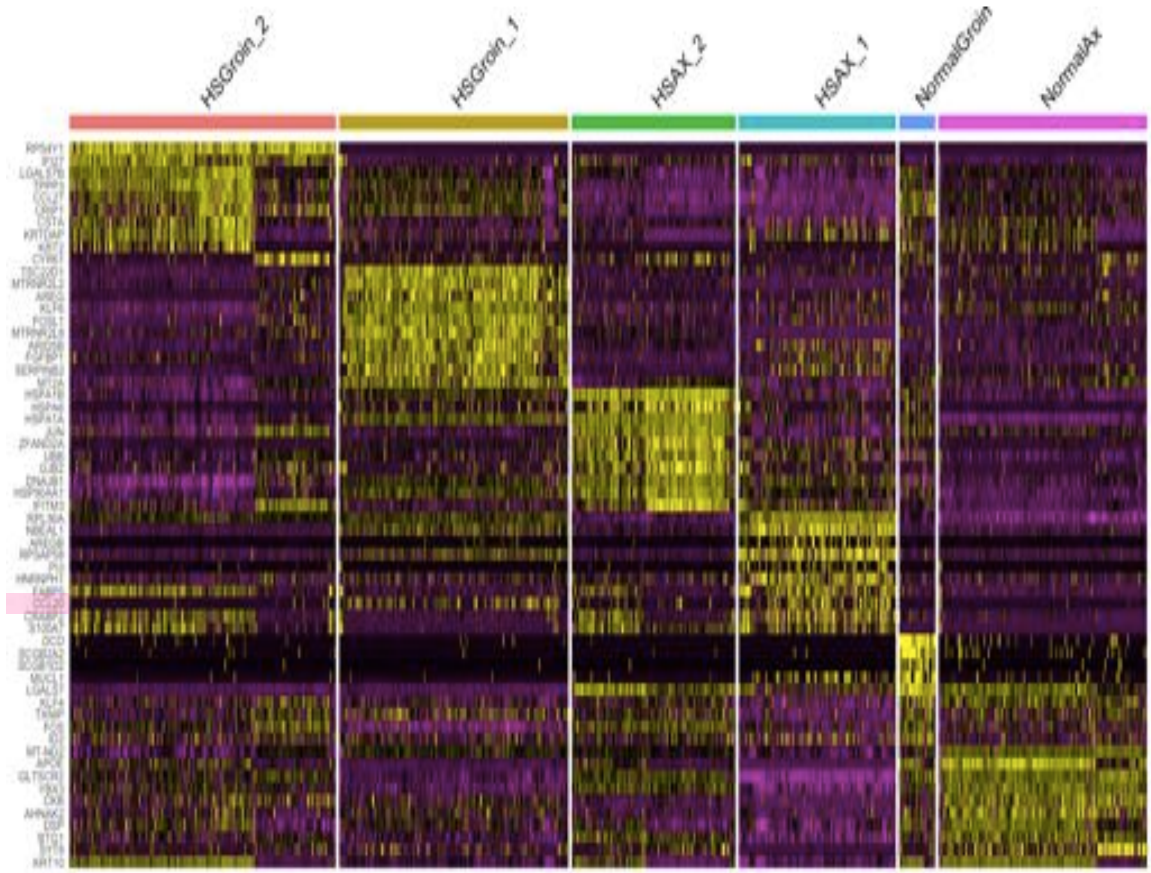

**Sup Figure 5. Heterogeneity in the surface epidermis between different patients.** (A) UMAP for epidermal clusters from different patients. (B) Heatmap showing the top unique genes expressed in different individuals.

### Sup Figure 6.

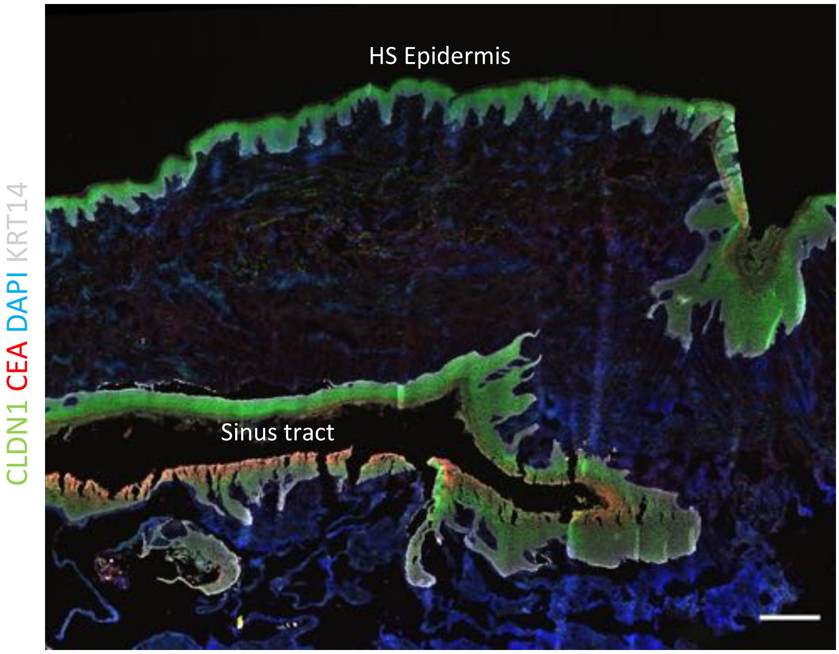

**Sup Figure 6. CLDN1 expression in HS epidermis and sinus tract.**
Immunofluorescent image, showing CLDN1 and CEA expression in HS lesional skin. Note that CLDN1 expression is significantly down-regulated in the sinus tract where CEA is also expressed. Scale bar, 500um.

**Sup Figure 7.**

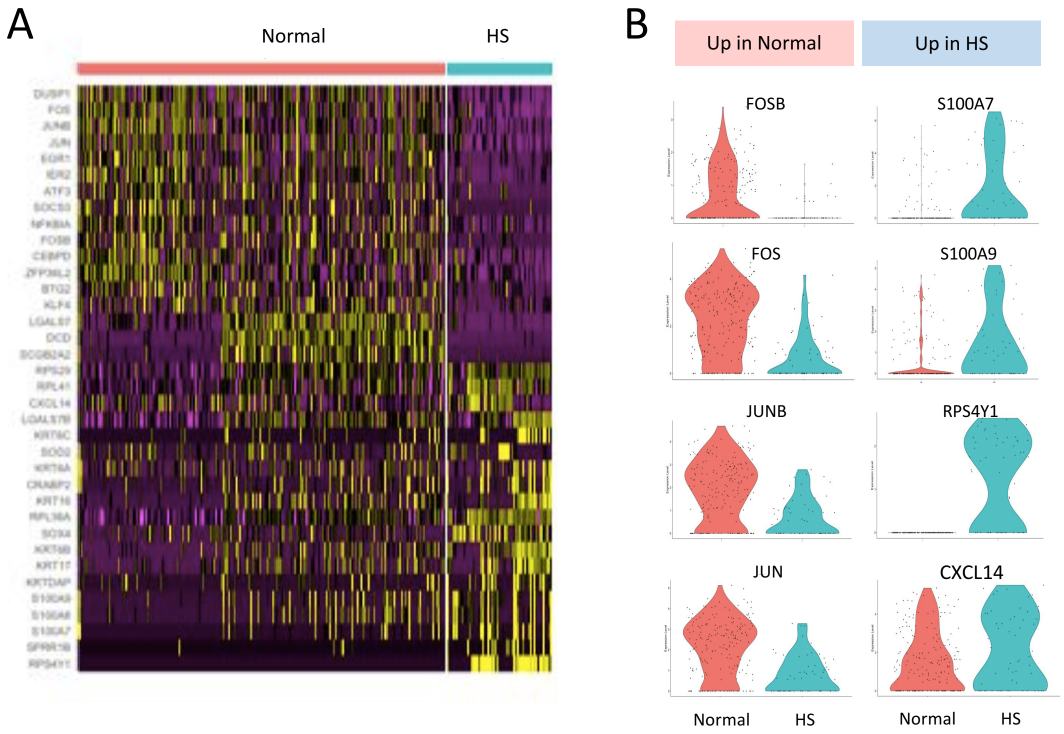

**Sup Figure 7.** scRNAseq analysis on hair follicle cluster. (A) Heatmap, showing top differentially expressed genes in normal and HS skin. (B) Violin plot, showing expression of genes in HF clusters that are changed between normal and HS skin.

**Sup Figure 8.**

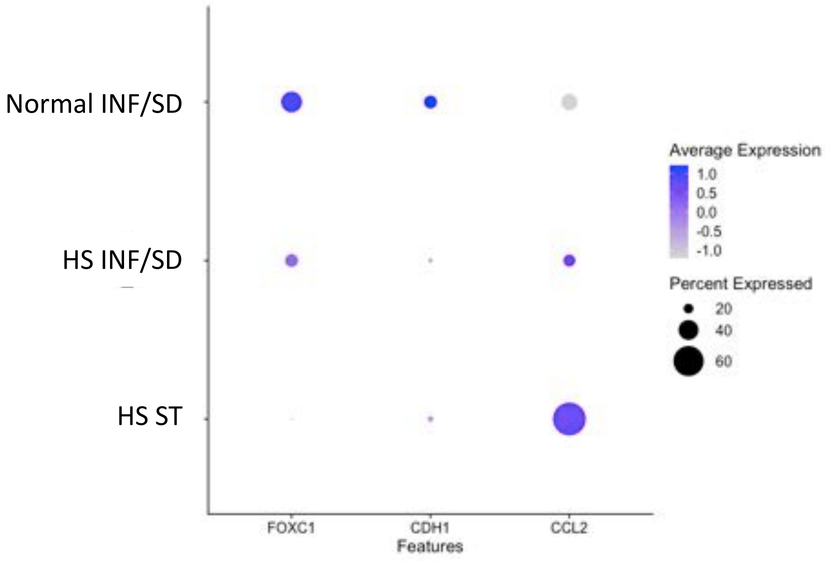

**Sup Figure 8.**
Dot plot, showing the average and percent expression of FOXC1, CDH1, CCL2 in normal, HS infundibulum/sweat duct (INF/SD), and HS sinus tract (ST)

**Sup Figure 9.**

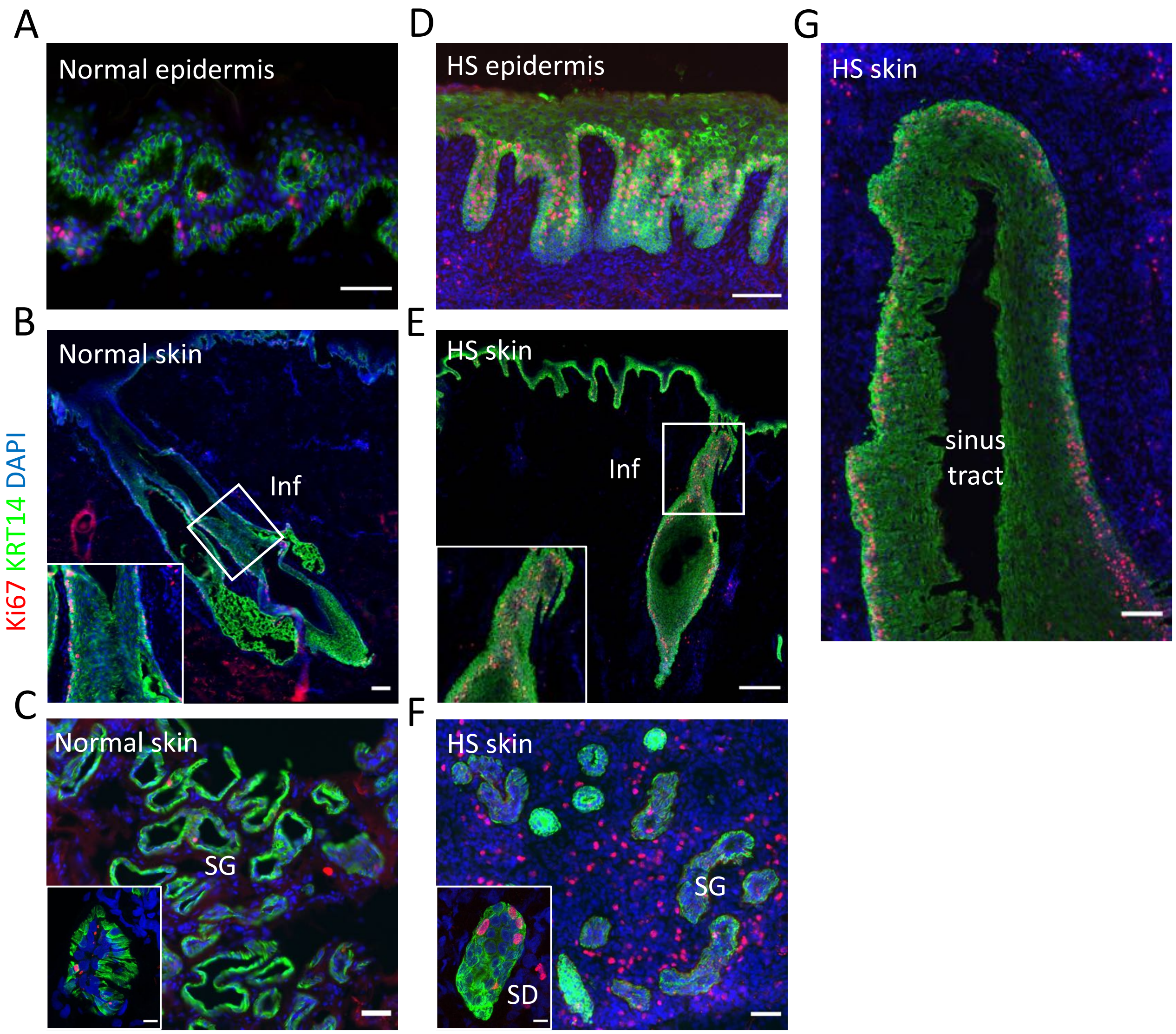

**Sup Figure 9. Cell proliferation in human normal and HS skin, appendages and sinus tract.** Immunofluorescent staining for Ki67 (red) in (A-C) normal skin and (D-G) HS lesional skin. Note that there are more proliferating cells in HS skin epidermis, infundibulum, sweat duct and sinus tract regions, but not sweat gland.. Inf, infundibulum; SG, sweat gland; SD, sweat gland. Scale bar, (A) 50 um, (B) 100 um, (C) 50 um/10 um (inbox), (D) 100 um, (E) 25 um, (F) 50 um/10 um (inbox), (G) 200 um.

**Sup Figure 10.**

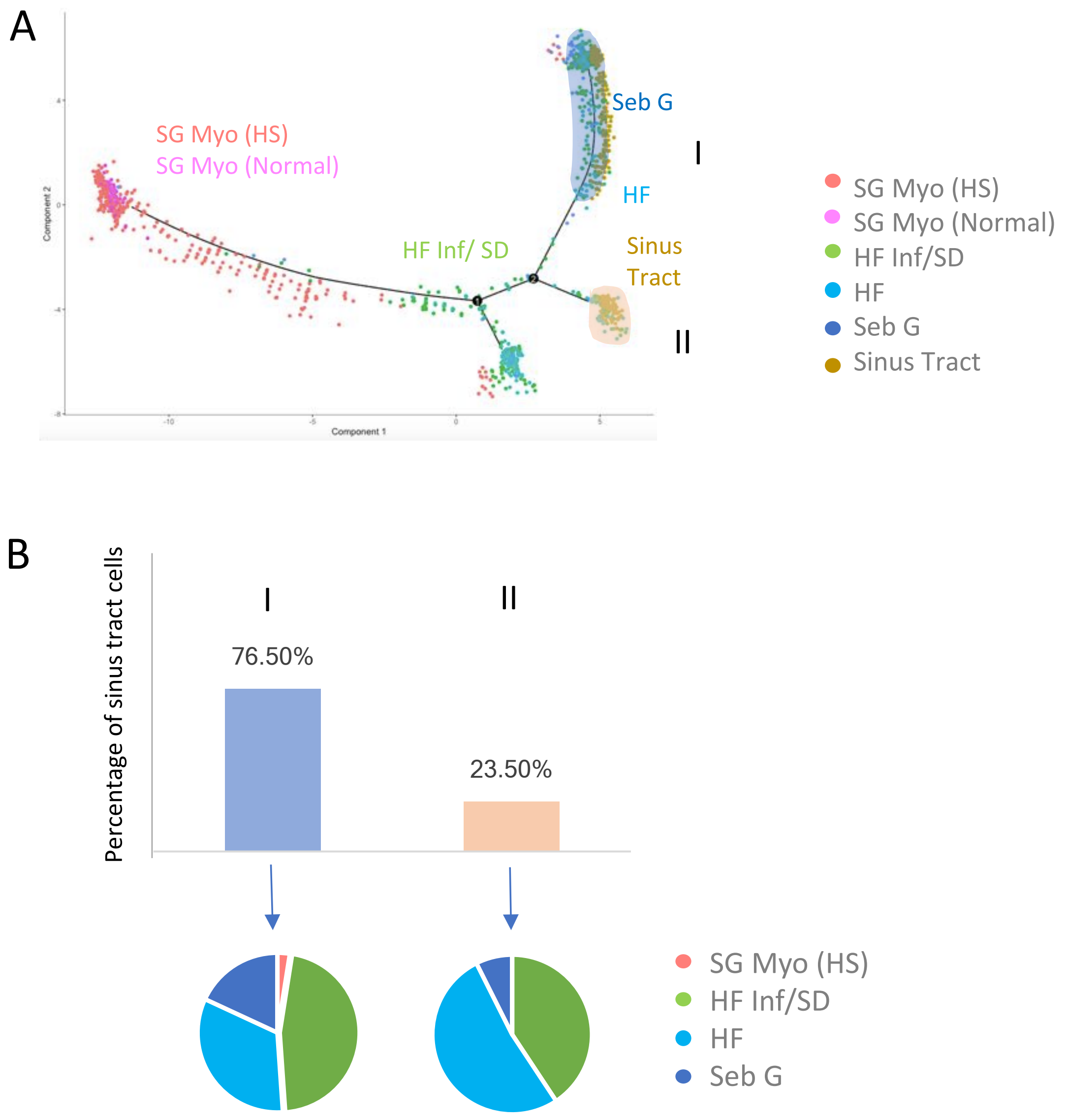

**Sup Figure 10.** (A) Pseudotime analysis, plot showing the cell trajectory of cell clusters from skin appendages and HS sinus tract. Note that cells from HS sinus tracts have a cell trajectory most similar to hair follicles and infundibulum/sweat duct. Blue-shaded area I and II, showing the cellular state where cells from HS sinus tract are distributed. (B) Bar graph, showing the percentage of cells from HS sinus tracts distributed in the state I and II, respectively. Lower pie chart showing the composition of the other cell types that share the same cellular state with the HS sinus tract cells. Note that in both state, majority of the cells are from Hair follicle infundibulum/sweat ducts.

**Sup Figure 11.**

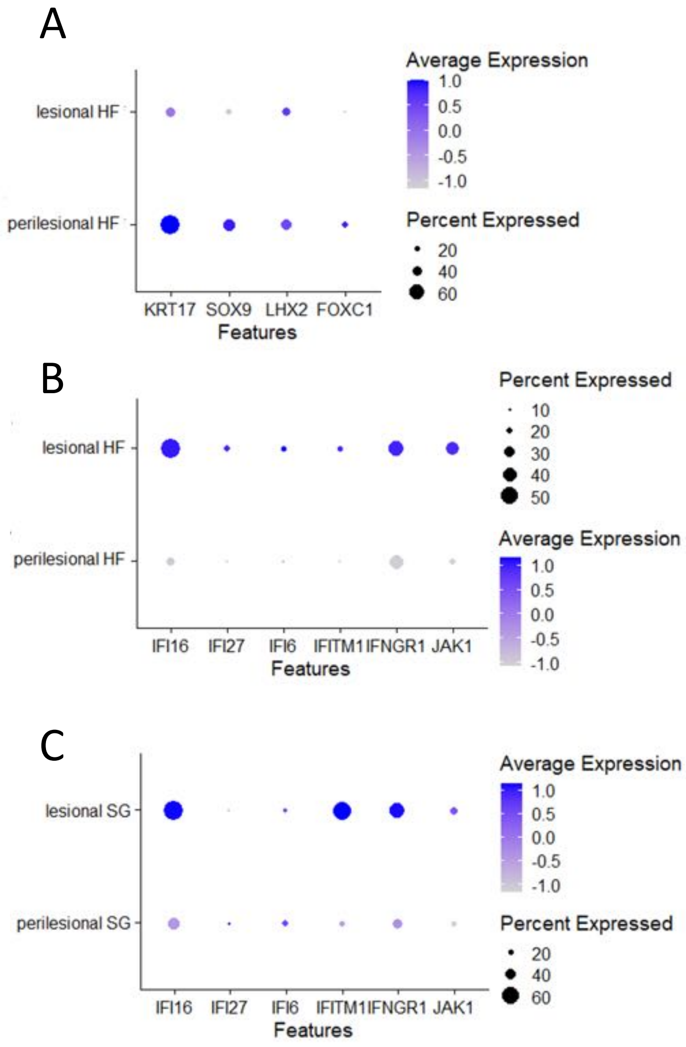

**Sup Figure 11.** Dot plots, showing (A) expression of HF stem cell marker genes in HF clusters, (B) expression of IFN signature genes in HF clusters, and (C) expression of IFN signature genes in SG clusters in HS skin samples.
